# Quantifying the evolution of SNPs that affect RNA secondary structure in *Arabidopsis thaliana* genes

**DOI:** 10.1101/2024.09.27.615253

**Authors:** Galen T. Martin, Christopher J. Fiscus, Brandon S. Gaut

**Author notes:** Author for correspondence: Brandon S. Gaut 321 Steinhaus Hall, University of California, Irvine Irvine, CA 92697.

## Abstract

Single-stranded RNA molecules can form intramolecular bonds between nucleotides to create secondary structures. These structures can have phenotypic effects, meaning mutations that alter secondary structure may be subject to natural selection. Here we examined the population genetics of these mutations within *Arabidopsis thaliana* genes. We began by identifying derived SNPs with the potential to alter secondary structures within coding regions, using a combination of computational prediction and empirical data analysis. We identified 8,469 such polymorphisms, representing a small portion (∼0.024%) of sites within transcribed genes. We examined nucleotide diversity and allele frequencies of these “pair-changing mutations” (pcM) in 1,001 *A. thaliana* genomes. The pcM SNPs at synonymous sites had an 13.4% reduction in nucleotide diversity relative to non-pcM SNPs at synonymous sites and were found at lower allele frequencies. We used demographic modeling to estimate selection coefficients, finding selection against pcMs in 5’ and 3’ untranslated regions. Previous work has shown that some pcMs affect gene expression in a temperature-dependent matter. We explored associations on a genome-wide scale, finding pcMs exist at higher population frequencies in colder environments, as do non-PCM alleles. Derived pcM mutations have a small but significant relationship to transcript abundance, however; alleles containing pcMs had an average reduction in expression of 137.4 normalized counts compared to genes with conserved ancestral secondary structure (mean expression = 3215.7 normalized counts). Overall, we document selection against derived pcMs in UTRs but with limited evidence for selection against derived pcMs at synonymous sites.

## INTRODUCTION

RNA molecules are single stranded (ssRNA), which gives them the ability to form Watson-Crick bonds between bases in the same molecule (Varani and McClain 2000). This intramolecular base pairing, termed secondary structure, largely determines the three-dimensional shape of the molecule. The capacity for an RNA sequence to form secondary structures affects the function of transcribed regions of genomes in many ways (Vandivier et al. 2016). For example, secondary structures influence function by modulating translation (Kozak 1988; Svitkin et al. 2001), mRNA splicing (Buratti and Baralle 2004), ribozyme activity (Steitz and Moore 2003), localization (Bullock et al. 2010), and protein-RNA interactions (Williams and Marzluff 1995). Additionally, they affect the epigenetic fate of genes by influencing their RNA stability (Li et al. 2012), complement of small interfering RNAs (siRNAs) and DNA methylation (Martin et al. 2022). The ultimate impact of a transcribed genomic region on phenotype (Duan et al. 2003) and fitness (Innan and Stephan 2001) is therefore shaped by its capacity to form secondary structures; for example, mutations that affect mRNA structure in humans have been implicated in disease (Halvorsen et al. 2010). Yet, the evolutionary dynamics of mutations affecting secondary structures in mRNAs have received little attention in the evolutionary biology literature, with most such studies focusing on non-coding RNAs (Nowick et al. 2019).

One interesting and unexplored aspect of selection on secondary structure is its potential to contribute to adaptation. In protein coding genes, positive selection can, in theory, act on both the amino acid sequence and the mRNA secondary structure. However, selection is typically considered through the lens of mutational effects on proteins. Metrics like d_N_/d_S_ (the ratio of nonsynonymous to synonymous substitution rates) are used to identify loci under positive or purifying selection, but this approach does not account for possible selection on mRNA secondary structure, which is likely to affect both the numerator and the denominator of dN/dS. Consider, too, that fitness optima of secondary structures may change with the environment. For example, Ferrero-Serrano et al. (2022) recently demonstrated that two experimentally-validated structure-changing SNPs [often termed “riboSNitches” (Halvorsen et al. 2010)] caused different folding dynamics in cold vs. warm environments. Thus, the fitness optima of secondary structures may vary with temperature and perhaps other environmental variables.

Selection on secondary structure could also have important methodological consequences for measuring selection with molecular data. This is because interpretation of d_N_/d_S_ ratios assumes that synonymous mutations are selectively neutral (Kimura 1968a). Since the 1980s, evolutionary biologists have known that this is not entirely correct because codon usage is non-random (Ikemura 1981), and more recent studies have demonstrated strong non-neutral fitness effects from synonymous mutations (Lawrie et al. 2013; Lebeuf-Taylor et al. 2019). With regard to the fitness effects of secondary structure, it is known that (i) secondary structures within mRNA coding regions are more stable than expected under randomized codon usage (Seffens and Digby 1999), (ii) the location of synonymous substitutions is not random with respect to secondary structure stability (Chamary and Hurst 2005), (iii) codon usage is constrained towards weaker structure around miRNA-binding sites (Gu et al. 2012), and (iv) synonymous variants disrupting computationally-predicted secondary structure exist at reduced frequencies in human populations (Gaither et al. 2021), implying that purifying selection acts on these variants. Approaches like d_N_/d_S_ and the McDonald-Kreitman test (McDonald and Kreitman 1991) typically rely on the assumption that synonymous changes 8are selectively neutral, but they have been shown to be sensitive to even weak selection (Rahman et al. 2021). Depending on the strength and prevalence of RNA-level selection, accounting for secondary structure could be important for distinguishing neutral synonymous variants from less neutral variants.

Another reason such mutations are evolutionarily interesting is through the possibility of different selective effects between the RNA and protein “life stages” of gene expression. Nonsynonymous mutations can alter both amino acid sequence and secondary structure stability, potentially leading to a conflict between selection for protein function (protein-level selection) and mRNA stability (RNA-level selection) (Wegler et al. 2020). For example, a derived missense substitution may enhance the effectiveness of a protein but have an overall deleterious effect by compromising mRNA fitness through a less favorable secondary structure, potentially leading to improper splicing, translation, or reduced stability (Vandivier et al. 2016). The frequency and importance of this potential pleiotropic antagonism depend on the relative strength of selection acting on mutations affecting secondary structure compared to mutations that affect amino acid sequence. If these conflicts exist, they will constrain the efficacy of positive selection (Fraïsse et al. 2019).

Finally, while secondary structures serve important functions, particularly strong secondary structures have unique properties that may negatively affect mRNA half-lives. Stable genic hairpins can cause genes to behave like pre-microRNA (miRNA) transcripts (Li et al. 2012), which form hairpin structures that are targeted by Dicer-like enzymes (Vergani-Junior et al. 2021) and are subsequently degraded into small RNAs. Like in pre-miRNA loci, elevated numbers of small interfering RNAs map to these structured genes (Li et al. 2012; Martin et al. 2023), putatively because their hairpins are degraded by Dicer-like enzymes. In turn, regions of miRNA-like secondary structure within genes correspond to high densities of small RNA mapping as well as high levels of small RNA-associated methylation (Martin et al., 2023), which often represses gene expression and function (Li et al. 2012). Given that small RNA mapping and repressive methylation are typically associated with silenced sequences, such as transposable elements, it is intriguing that many genes (up to 70% of annotated Zea mays genes) contain hairpin secondary structures (Martin et al. 2023). It is possible that these regions represent evolutionary conflicts between crucial secondary structure and downstream epigenetic effects.

In this study, we focus on the population genetics of mutations that are likely to affect secondary structure within the genes of the *Arabidopsis thaliana* (Arabidopsis) 1001 Genomes dataset (1001 Genomes Consortium 2016). We begin by establishing a method to identify SNP variants that are within the ancestral secondary structures of expressed coding regions. There are generally two approaches to identify these structures, and neither is perfect. The first is empirical measurement. X-ray crystallography can accurately determine the structure of a transcript (Zhang and Ferré-D’Amaré 2014), but it is prohibitively expensive and infeasible to perform on a genome-wide level. Sequencing approaches such as double-stranded RNA (dsRNA) sequencing (Zheng et al. 2010; Li et al. 2012), structure-seq (Ding et al. 2014), and SHAPE-seq (Kwok et al. 2013; Liu et al. 2021) have also been used to estimate secondary structures. These methods can be error prone, depend on coverage, and capture only a single possible secondary structure in a moment in time. The second approach is computational prediction [e.g., (Halvorsen et al. 2010; Lorenz et al. 2011; Zhang et al. 2020)], which is widely used but does not always recapitulate known structures from X-ray crystallography (Zhang et al. 2020). Nonetheless, newer prediction methods have become more accurate and have distinct advantages, such as the ability to integrate information across many possible secondary structures (Zhang et al. 2020).

Here we adopt a combined approach that uses both computational prediction and sequencing data to identify SNPs that may affect secondary structures in expressed coding region of *A. thaliana*. We then calculate the population frequencies of these variants across the 1001 global Arabidopsis accessions to address four sets of questions. First, is there evidence that these mutations are under selection? That is, do they have evolutionary histories similar or dissimilar to putatively neutral synonymous mutations? Second, if they appear to be under selection, what is the inferred strength of selection compared to synonymous and nonsynonymous SNPs that are not within secondary structures? Third, is there any evidence to suggest that RNA-level selection conflicts with protein-level selection or that they affect one possible phenotype, gene expression? Finally, previous work has suggested that secondary-structure altering SNPs may be associated with environment variables, particularly temperature. Is this true on a genome-wide scale?

## RESULTS

### Identifying unpaired mutations (upM) and pair changing mutations (pcM) mutations

Identifying causative mutations that change the overall conformation of an RNA molecule is a complex and unsolved problem (Ferrero-Serrano et al. 2022). We developed a method to identify derived mutations with a high likelihood of being ancestrally paired (that is, hydrogen bonded to another nucleotide base in the ancestral *A. thaliana* genome) within secondary structures. We refer to such mutations as “pair changing mutations (pcM)”, while those that are not ancestrally paired are “unpaired mutations (upM).” To classify these variants, we followed three steps (Figure 1). First, we polarized ancestral SNPs from the Arabidopsis 1001 Genomes Project (1001 Genomes Consortium 2016) using an A. lyrata outgroup (see Methods). We used these ancestral SNPs to create an ancestral pseudo-transcriptome from the TAIR10 assembly (Berardini et al. 2015) by replacing derived alleles present in the Col-0 reference genome with the inferred ancestral SNP. Second, we extracted mRNA sequences from the pseudo-ancestral reference and inferred base-pairing potentials within these sequences using LinearPartition (Zhang et al. 2020). LinearPartition calculates a partition function for a complete RNA sequence, and it sums equilibrium constants for all possible secondary structures for a sequence (i.e, not just the most likely structure). It outputs a base-pairing matrix that conveys the estimated probability that two bases pair. We focused only on bases with a high (>0.90) probability of pairing. Finally, we overlapped LinearPartition analyses with empirical data, namely previously-generated dsRNA data (Zheng et al. 2010). The data were generated from 6-week-old Col-0 (flower bud clusters, leaves, and all aerial portions), and we used the data with the intention of distinguishing true paired bases from less-likely paired bases. In the sequencing data, we found that dsRNA regions were, on average, 26.9 nt long, and spanned a total of 1.9 × 10 nt, representing 1.7% of the total mRNA database.

**Figure 1.**
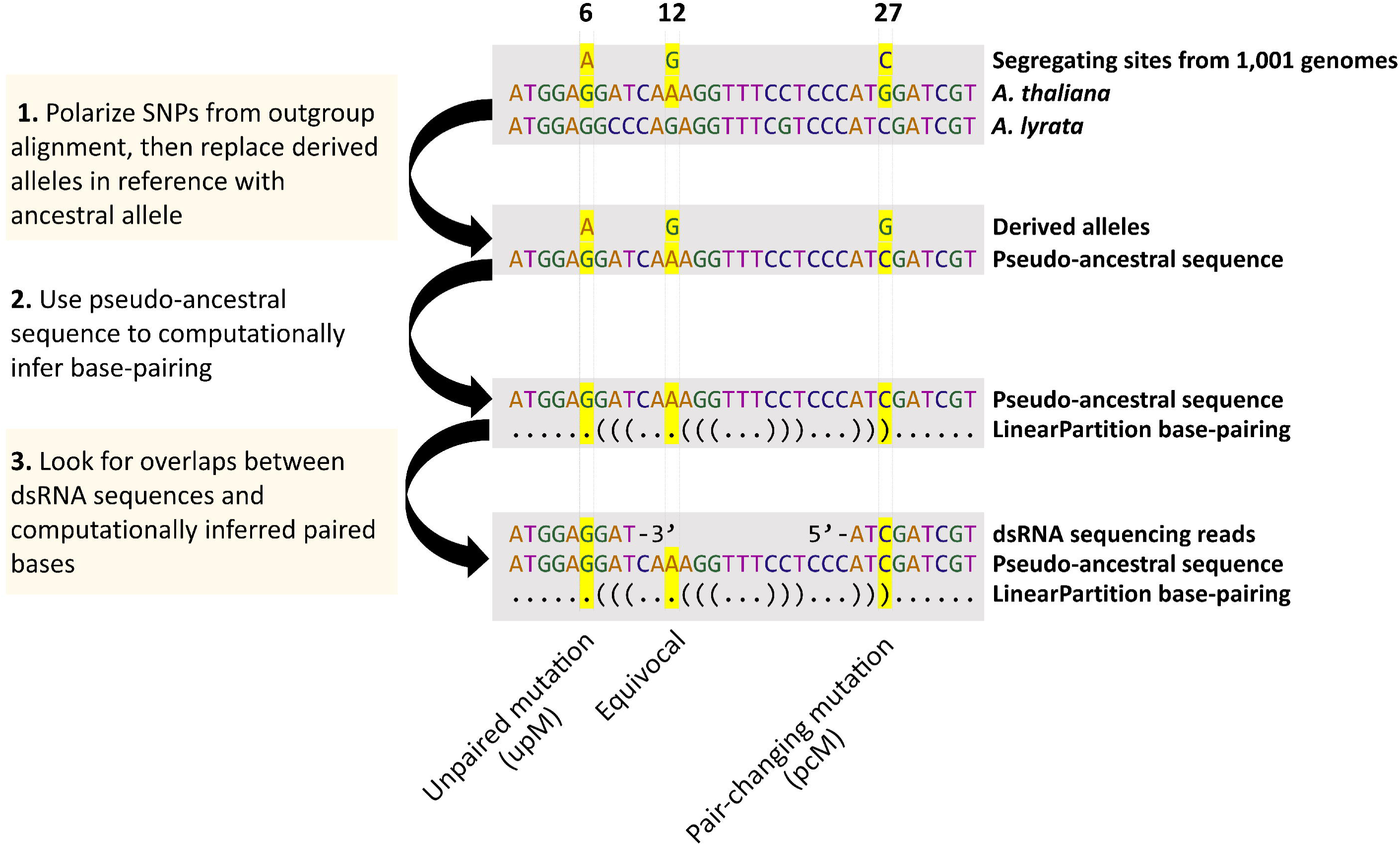
A schematic representation of the method used to identify derived unpaired (upM) and pair-changing (pcM) mutations. Three SNPs are shown at positions 6, 12, and 27 within a hypothetical gene: one that is unpaired (6, left), one that is equivocal (position 12, middle) and one that is a pair-changing mutation (position 27, right). Bases with high (>0.90) LinearPartition base-pairing probabilities are shown as parentheses. Both dsRNA and LinearPartition evidence were required to designate a base as pcM; bases with only one type of supporting evidence (either base-pairing probabilities > 0.90 or empirical dsRNA evidence) were deemed equivocal.

Given both LinearPartition and dsRNA data, we defined derived pcM mutations as the subset of SNPs: i) that had a LinearPartition probability >0.9, ii) that were detected as paired in dsRNA data, and iii) whose presumed paired base did not also contain a complementary SNP. For example, if the identified base contained an A to G mutation at position 1 and was found to be ancestrally paired with a T at position 20, then the SNP at position 1 was not counted if position 20 contained a T to C substitution. This approach yielded three sets of derived SNPs: i) pair-changing mutation (pcM), which were predicted to alter base-pairing based on both empirical and computational evidence, ii) ‘equivocal’ SNPs that were predicted to alter base-pairing based on only one of the two prediction methods, and iii) unpaired mutations (upM) that had no evidence for being within secondary structures. After applying these rules, the pcM set consisted of 8,469 inferred mutations across 5,141 Arabidopsis genes (Table 1), representing a subset of 201,965 ancestrally paired bases (Table 1). Note, however, that the pcM SNPs likely do not reflect all of the bases involved in secondary structures, due to inevitable false negatives in our conservative approach. To address this concern, we also compared results from less conservative pcM/upM definitions (e.g., changing the pairing-probability cutoff and relying only on LinearPartition by not considering dsRNA overlap) to the pcM sets. The less conservative datasets yielded quantitatively similar downstream results (see below), and so for simplicity we focus primarily on analyses with the pcM set.

**Table 1.** Genomic sites categorized by effect inferred effects on secondary structure.

|  | Ancestrally paired | Ancestrally unpaired | Equivocal<sup>1</sup> |
| --- | --- | --- | --- |
| Total number of bases | 201,965 | 63,158,076 | 1,305,821 |
| Total number of SNPs | 8,469<sup>2</sup> | 2,320,555<sup>3</sup> | 70,719 |
| Nonsynonymous SNPs | 3,790 | 864,006 | 22,231 |
| Synonymous SNPs | 3,214 | 631,838 | 16,038 |
| UTRs (5' + 3') | 1,103 | 438,206 | 240,526 |

<sup>1</sup>Supported as paired by the computational method (LinearPartition) or by dsRNA coverage but not both.
<sup>2</sup>These SNPs represent the total set of pair-changing mutations (pcM).
<sup>3</sup>These SNPs represent the set of unpaired mutations (upM).

### Prevalence and distribution of upM and pcM mutations

If mutations that affect secondary structure are selectively important, one naive expectation is that their distribution across the genome differs from those mutations that do not affect structure. To profile and compare distributions among SNP types, we categorized derived SNPs based on their predicted impact based on SnpEff annotations (Cingolani et al. 2012). upMs were more likely to occur in untranslated regions (UTRs) and splice sites than pcMs, which were more common in coding regions (Table 2). pcMs were about half as likely to be found in the UTRs as expected under random distributions based on the length of features within genes—i.e., 3’ UTRs comprised ∼15% of genic space but only 8.04% of pcMs were within UTRs. Similarly, the percentage of 5’ UTR pcM mutations (4.98%) was lower than expected given the percentage of paired bases within 5’ UTRs (6.86%).

**Table 2.**
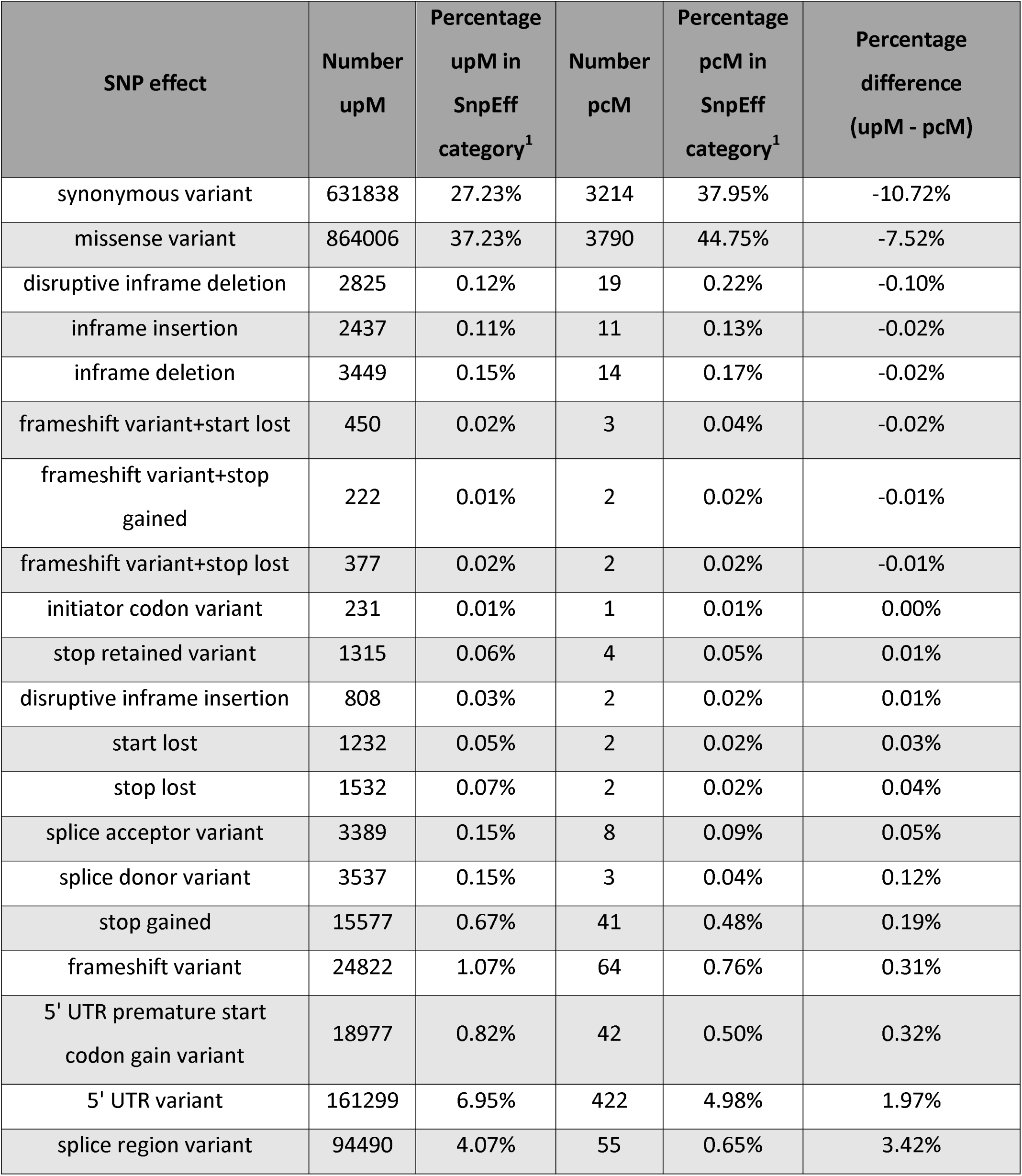
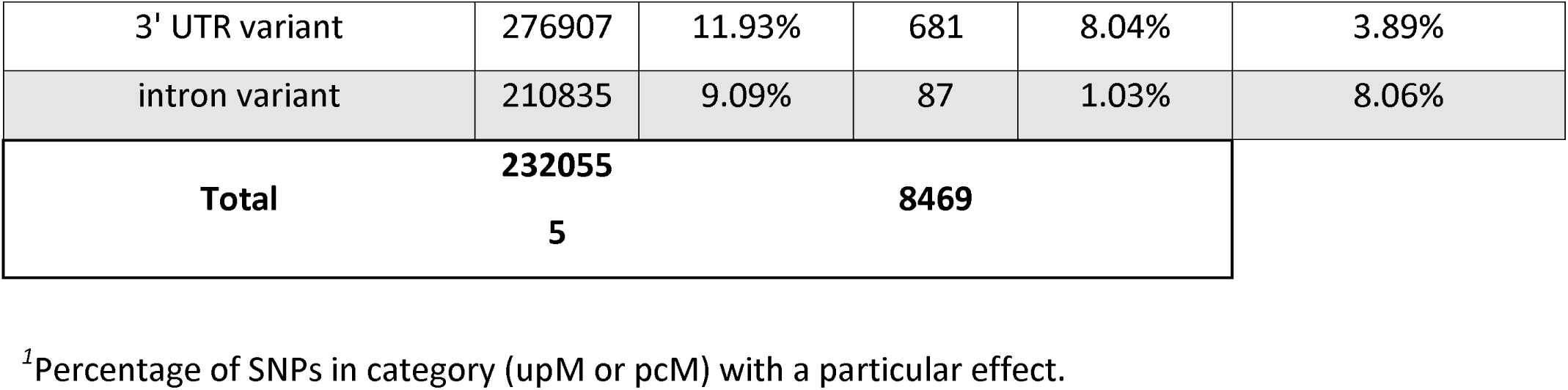
SnpEff annotations for upM vs pcM SNPs.

These observations could be caused either because secondary structures are less frequent in UTRs or because selection against pcM mutations is stronger in UTRs. To explore these options, we compared the locations of 8,469 pcMs to the set of 201,965 ancestrally paired bases that did not have a derived SNP. If selection is not the cause, we reasoned that the distribution of pcMs in UTRs should be similar in proportion to ancestrally paired bases. To perform each permutation, we chose a random subset of n = 8,469 pseudo-pcM sites from the complete paired site dataset of n= 201,965. For each permutation, we counted the percentage of randomly-assigned sites in each genic component (5’ UTR, coding region, and 3’ UTR) and then compared those proportions to observed values. We found, for example, that the observed proportion of 13% of pcMs in 5’+3’ UTRs differed significantly (P < 0.01) from the 16% proportion of paired ancestral sites in UTRs. These results suggest that the locational skew in pcMs may not be due solely to locational biases but may be consistent with selection shaping the location and distribution of pcMs.

### Reduced nucleotide diversity at paired versus unpaired sites

Nucleotide diversity at synonymous sites (π_S_) tends to be higher than at nonsynonymous sites (π_N_), which is generally interpreted as the result of purifying selection (Ingvarsson 2010; Osada 2015). We calculated nucleotide diversity on different sets of segregating sites. For pair-changing mutations, we focused only on synonymous sites (syn_pcM) to avoid the confounding effects of selection on nonsynonymous substitutions. We compared nucleotide diversity at segregating synonymous sites between the syn_pcM (n=3,214) and syn_upM (n=631,838) categories. We hypothesized that, if syn_pcM mutations are neutral, syn_pcM diversity (π_syn_pcM_) should be equivalent to syn_upM diversity (π_syn_upM_). Alternatively, if selection on paired bases is strong and similar to that of protein-level selection, π_syn_pcM_ should be similar to nonsynonymous diversity (π_N_) (n=864,006). We found that π_syn_pcM_ (median: π_syn_pcM_ = 0.0071, mean: πsyn_pcM = 0.071) was significantly lower than π_syn-upM_ (median: π_syn_upM_ = 0.0082, mean: π_syn_upM_ = 7.8e-2) (Figure 2A; t-test P < 0.001). However, π_syn_pcM_ was also significantly higher than π_N_ (median: π_N_ = 0.0037; mean: π_N_ = 0.050), putting pcM diversity at an intermediate level (Figure 2A; t-test P < 0.001). These results hint at purifying selection on syn_pcM sites compared to syn_upM sites.

**Figure 2.**
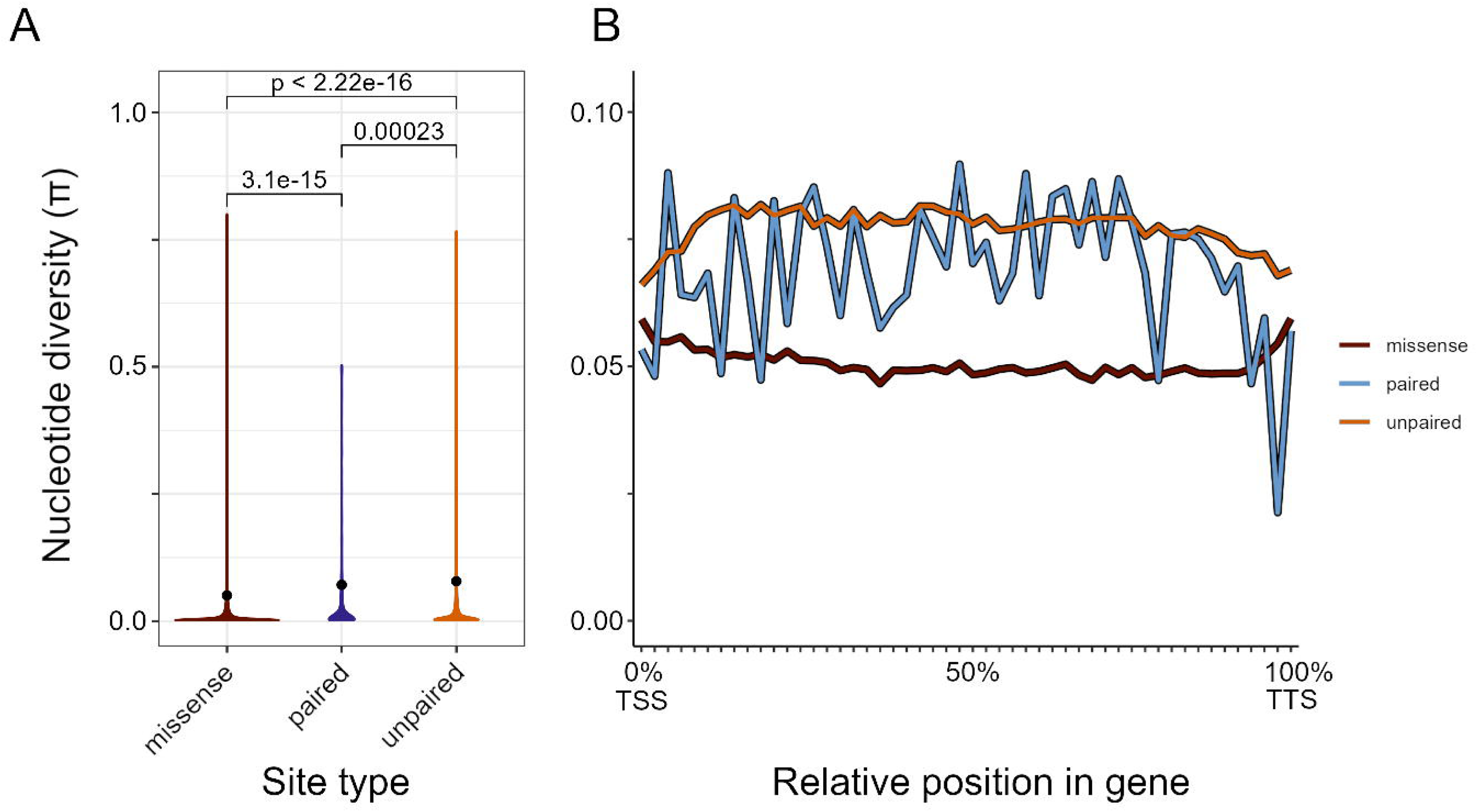
Nucleotide diversity at synonymous paired (pcM), synonymous unpaired (upM), and nonsynonymous (missense) sites. (A) Violin plot of nucleotide diversity at different site types, with the black dots representing mean diversity. Statistical contrasts are provided above the violin plots, based on t-tests. (B) π calculated in windows across the length across all analyzed CDS regions. The x-axis represents length-standardized windows across the span of all analyzed genes from the 5’ end (transcription start site, TSS) to the 3’ end (transcription termination site. TTS). Missense refers to nonsynonymous bases not identified as pair-changing mutations; paired refers to synonymous pcM mutations; unpaired refers to synonymous upM mutations.

An open question is whether selection on secondary structure interferes with selection for amino acid sequence. One factor that may influence this relationship is variation in selection across the length of genes. For example, secondary structure is known to be particularly important at start codons and intron splice sites (Li et al. 2012; Vandivier et al. 2016), while the amino acid sequence is more important towards the middle of the protein (Bricout et al. 2023). To explore spatial distributions, we measured π at the three different types of sites across the length of gene coding sequences (Figure 2B). The distributions differed visually among site categories. π_N_ was lowest at the middle of the coding sequence, while syn_π_upM_ was lowest towards the edges. The signal for π_syn_pcM_ was noisy, likely owing to the low n of this category, but it dipped towards the 3’ end of the coding sequence.

### Investigating selection on structural mutations

Given results based on π, we predicted that mutations that putatively change secondary structure are generally more deleterious than synonymous changes that do not affect secondary structure. To examine this prediction more formally, we calculated the site frequency spectra (SFS) of various classes of mutations and then used the SFS to infer the strength of selection. The distribution for pcM sites was skewed visually towards lower frequency alleles compared to upM sites for both synonymous and nonsynonymous sites (Figure 3AB), reflecting more singletons and fewer intermediate and fixed mutations. The set of pcM mutations was a small subset of total SNPs, so we tested for statistical significance between SFSs in two ways: a Kolgorov-Smirnov test and a permutation-based approach (Figure 3CD). The SFS for pcM mutations at both synonymous and nonsynonymous sites differed significantly from the SFS of syn_upM mutations. Nonsynonymous pcM (i.e., non_pcM) mutations had a stronger skew toward rare variants than syn_pcM mutations (Figure 3AB), with a correspondingly lower significance value relative to the syn_upM SFS (permutation non_pcM, P ≃ 0; syn_pcM, P = 0.01) (Figure 3CD). Finally, we note that the SFS of mutations in the equivocal class (i.e., those which were identified by either the computational or sequencing approach, but not both) fell between the pcM and upM distributions (Figure 3AB).

**Figure 3.**
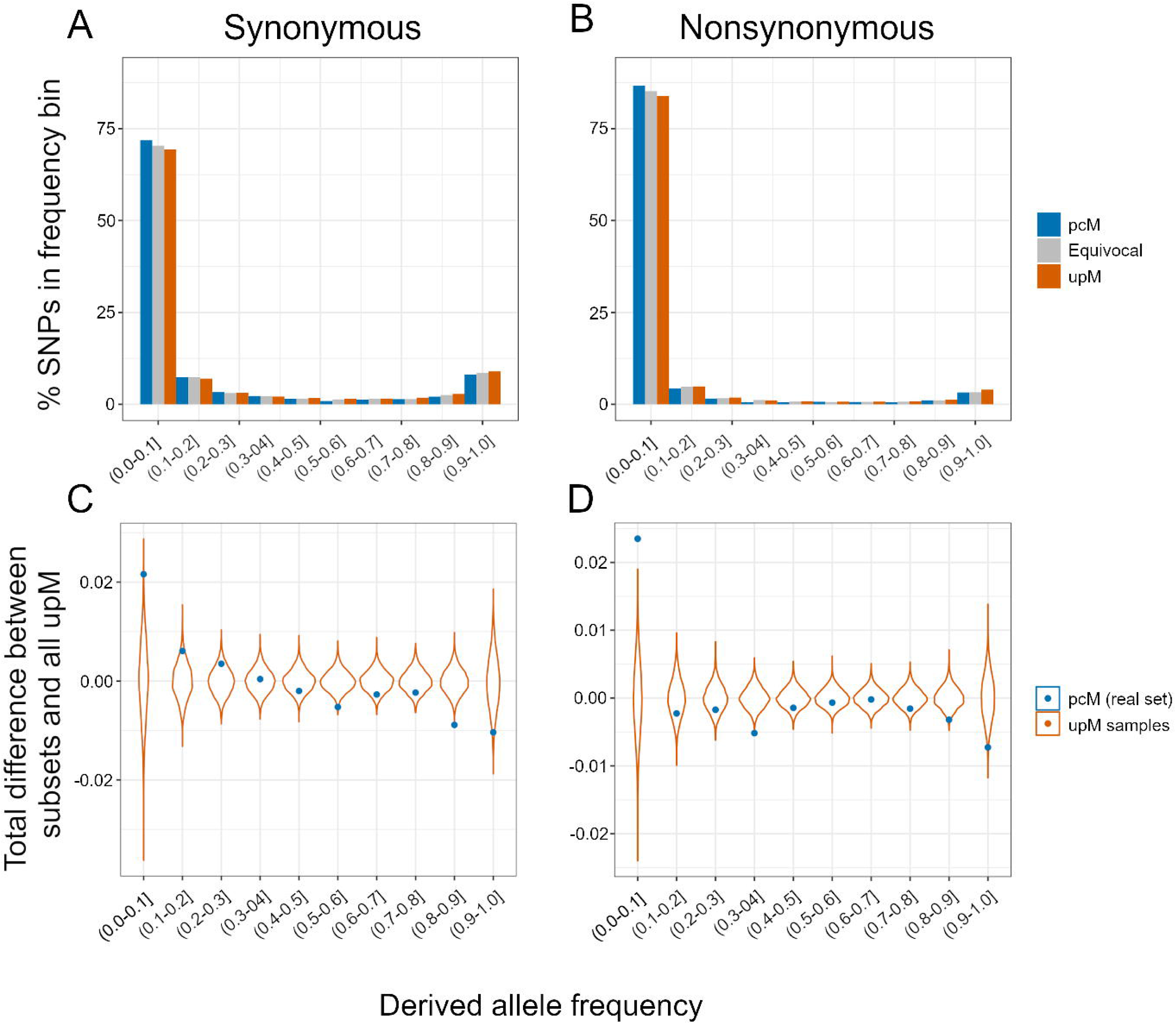
(A) Unfolded site frequency spectra (SFS) showing derived allele frequencies of synonymous alleles categorized by their inferred effect on ancestral secondary structure. (B) SFS of nonsynonymous alleles in each category. (C) Permutation distributions for differences between paired and unpaired SFS at synonymous sites (black dot shows missense for scale) and (D) for nonsynonymous sites. In panels C and D, violins represent the distribution of differences in random samples (same n as paired sites) from the unpaired data, while points show the true differences. Differences (y-axis) were calculated as the percentage of alleles in each pcM SFS bin subtracted from the percentage in the same upM SFS bin (e.g., {0-0.1] in pCM minus {0-0.1] in upM etc.)

We evaluated the robustness of these results by investigating datasets based on alternative definitions of pcM and upM. First, we considered sites identified as likely to be paired by LinearPartition, without filtering by dsRNA overlap. For both synonymous (n = 19,252) and nonsynonymous (26,021) pcM, allele frequencies remained significantly different compared to syn_upM (Figure S1). Second, we loosened cutoffs for base-pairing probabilities inferred by LinearPartition, while continuing to filter by dsRNA cutoff. With a base-pairing probability threshold of 0.50 (opposed to the original 0.90), the SFS of both syn_pcM (n = 20,596) and non_pcM (n=57,349), the SFSs remained significantly different to the SFS of syn_upM (Figure S2).

We used SFS information to infer the strength of selection on pcM mutations using fitDadi (Gutenkunst et al. 2009; Kim et al. 2017), which estimates demographic history from frequency spectra of neutral alleles. Here we used syn_upM mutations for demographic inference, reasoning that they represent the most likely set of neutral sites in our dataset. We fit four models with syn_upM SNPS, including a standard neutral model, an exponential growth model, a bottleneck model and a two-epoch model (Figure S3) (Methods). Based on the Akaike Information Criterion (AIC), the exponential growth model best fit the data (Table 3), with exponential growth starting 0.243 2N_Ancestral_ generations before present to a contemporary population size nu = 2.26 X N_Ancestral._ Similar shifts in population size were inferred with different demographic models, all of which inferred a ∼2-fold increase in population size beginning ∼0.4 N_Ancestral_ generations ago (Table S1). These inferences match previous work on *A. thaliana*, which is thought to have expanded from refugia after the last glacial maximum ∼20 KYA (François et al. 2008; Durvasula et al., 2017).

**Table 3.**
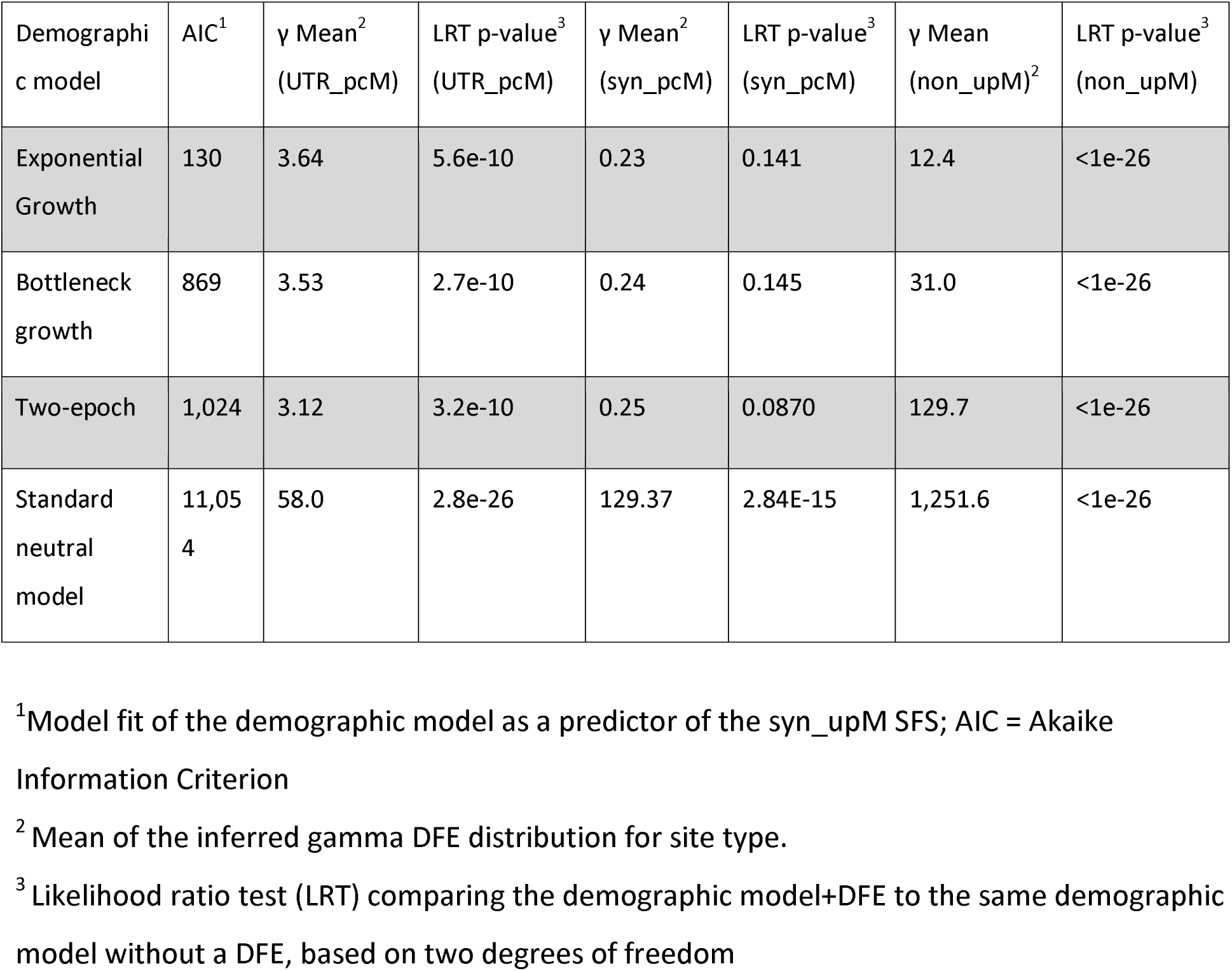
Demographic models used in fitDadi DFE estimation with information about inferred DFEs from various site categories.

We then used the fitted demographic models to estimate distributions of fitness effects (DFEs) from the SFS of syn_pcMs, pcMs in 5’+3’ UTR regions (UTR_pcMs) and non_upM SNPs. Based on visual analysis of the SFS (Figure 3AB), we expected the DFE to be weaker for syn_pcMs than for non_upMs, and this was the case (Figure 4A). From the inferred DFE based on the exponential growth model, syn_pcM sites had a mean scaled selection coefficient (γ = 2N_Ancestral_S) = 0.23, which was ∼50-fold smaller than the mean effect for non_upMs (Table 3). [Here, higher mean γ values denote stronger selection against derived mutations.] These estimates of DFE rely on the accuracy of the demographic model used, so we also compared DFEs estimated using the other demographic models, finding similar estimates for the magnitude of pcM selective effects (Table 3). Interestingly, the inferred mean scaled selection coefficient for UTR_pcMs was ∼15 higher than for syn_pcMs under the exponential growth and bottleneck models (Table 3; Figure 4B), reflecting the strong-left leaning skew for the UTR_pcM SFS (Figure S4). Similar estimates were obtained when 5’ and 3’ UTRs were examined separately (exponential growth model: 5’ UTR_pcM mean γ = 2.86; 3’ UTR_pcM mean γ = 5.87). Finally, we note that the relatively high mean values were not typical of UTRs, because non-pair changing mutations in UTRs had much lower mean γ values (exponential growth model: 5’ UTR upM mean γ = 0.11; growth: 3’ UTR upM mean γ = 4.0e-4)(Figure 4B).

**Figure 4.**
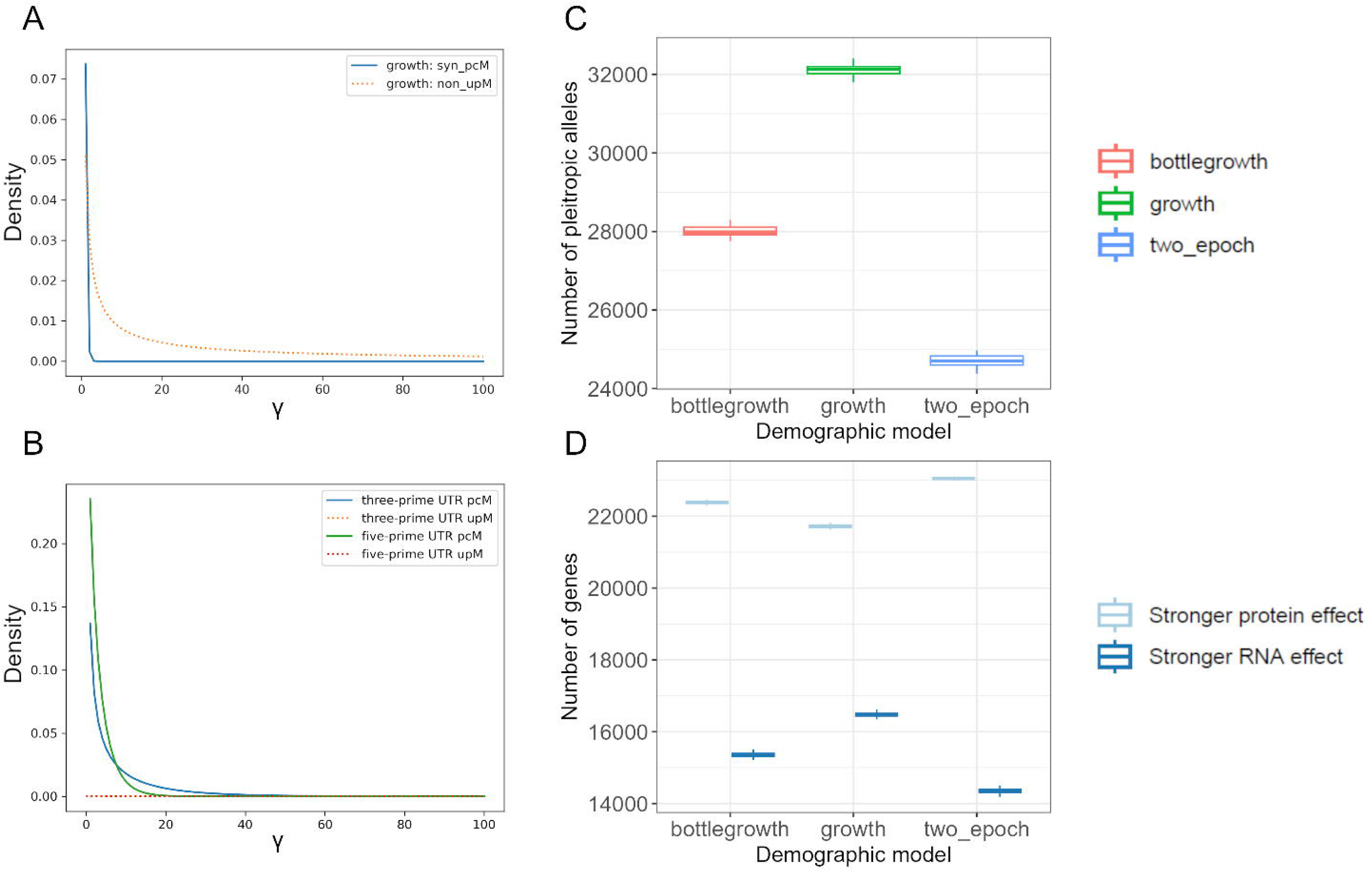
The inferred distribution of fitness effects (DFE) for mutation types. (A) The gamma DFE distribution for syn_pcM sites and non_upM sites. The x-axis is the scaled selection coefficient, γ (=2N_ancestral_s); higher values on the x-axis refer to stronger purifying selection. Note that scale parameters for the syn_pcM distribution were not significantly different from zero (Table 3). (B) The gamma DFE distribution inferred from mutations within UTRs. pcM mutations from within 5’ (fiveprime) and 3’ (threeprime) regions are shown separately, as are upM mutations on the dotted lines. The axis are described in panel A. (C) The results of the pleiotropy simulations showing the number of sites within a genome where a simulated nonsynonymous mutation experienced stronger selective effects at the RNA (secondary structure) level than at the protein level. The numbers differ markedly when DFEs from different demographic models (exponential growth, bottleneck with growth and two epoch models) were used; however, as indicated on the y-axis, for each model > 24,000 nonsynonymous sites across the genome were expected to have stronger selection on secondary structure than a missense change. (D) The number of genes expected to have at least one site that has a stronger effect on protein (i.e., missense) change than the RNA (i.e., change in secondary structure) and vice versa. The boxplots in panels C and D refer to distributions of simulated values across 100 different simulations for each demographic model; within boxplots, the line shows the median value and the box edges show the interquartile range.

Since γ values < 1 are typically considered neutral, the mean DFE estimates suggest that syn_pcMs (mean γ = 0.23) do not have strong selective effects on average. We tested this idea more formally using likelihood ratio tests (LRTs) that compared nested models with and without DFEs. While the results did depend on the demographic model, the null hypothesis of γ shape and scale parameters equalling 0.0 could not be rejected under the exponential growth and bottleneck model (p = 014; Table 3). By contrast, the null hypothesis could be rejected for UTR_pcMs and for non_upM SNPs across all of the considered demographic models (LRT P-values < 1e-5) (Table 3) (but not for 5’ or 3’ UTR_upM mutations; P-values > 0.2) Altogether the DFE results detect selection against derived nonsynonymous SNPs and pair-changing mutations in UTRs, without strong statistical evidence for selection against syn_pcMs SNP.

### The potential for pleiotropy between secondary structure and amino acid changes

The DFE analyses suggest that derived syn_pcMs are effectively neutral on average, but nucleotide diversity and the SFS suggest the possibility of purifying selection against syn_pcMs (e.g., Figure 2A, 3CD). In any case, the inferred syn_pcM DFE provides a sense of the magnitude and variation of effects across synonymous sites within protein coding regions due to secondary structure alone. In contrast, the DFE based on non_upM mutations provides insights into selection pressures on amino acid changes. Together these two DFEs provide an opportunity to assess how often selection on RNA secondary structures is strong enough to interfere with selection on protein function at individual nucleotide sites.

To contrast RNA-level vs. protein-level selection, we performed simple simulations of selection effects within gene coding regions. The simulations consisted of four steps (Figure S5 provides a schematic). First, we began with 24,820 genes representing the length of genes that have Ensembl-based A. lyrata orthologs (mean length = 1275.4 nt). Second, for each gene we made the common (e.g., Kimura 1968b) but simplifying assumption that mutations at ∼76% of sites represent non-synonymous changes. Third, since our empirical analyses indicated that 0.43% of nonsynonymous alleles were also pcM mutations (Table 1), we randomly assigned this proportion of nonsynonymous mutations to affect secondary structure. At these nonsynonymous sites, we assigned two fitness effects: a “protein-level” fitness effect (γ_protein_) and a “RNA-level” fitness effect (γ_structure_). The two fitness values were assigned by drawing from the corresponding non_upM and syn_pcM gamma DFE distributions, as inferred from the exponential growth model. Finally, we tallied metrics from each simulation, such as the number of sites where RNA-level selection, as reflected by γ_structure_, was larger than γ_protein,_ and also the number of genes where this occurred for at least one site.

Using the DFEs inferred from the exponential growth model, we estimated that there were, on average, 97,268 sites across 23,190 genes (85% of genes) where the selection coefficient for the amino acid change was higher than that for a pcM mutation. In contrast, a much smaller but still substantial number of 22,118 sites encompassed the opposite case, where γ_structure_ > γ_protein_. These sites were found in 49% (or 13,437) of genes (Figure 4BC). While these estimates are subject to numerous caveats (see Discussion), they suggest that even the small fitness effects observed among syn_pcMs could interfere with protein-level selection at individual sites across a large subset of genes.

### Testing for expression and temperature effects for pcMs

Previous work on experimentally-validated secondary-structure polymorphisms have suggested that the affected gene expression in a temperature dependent matter temperature (Su et al, 2018). This observation, coupled with previous observations that gene expression varies with the presence and strength of secondary structures (Li et al., 2012; Vandivier et al., 2016; Martin et al., 2023), prompted us to test two hypotheses. The first is that the presence of pcMs correlate with gene expression. To investigate the potential for this effect on a genome-wide level, we used the expression data from the 1001 Genomes dataset and constructed a linear model with mixed effects similar to Muyle et al. (Muyle et al. 2021) (see Methods). The model measured within-gene expression differences between alleles and tested for the significance of the allelic state (pcM vs. no pcM) across all genes, ignoring genes without pcM SNPs. Genome-wide, our model detected that derived pcM alleles had significantly lower levels of expression compared to non-pcM alleles (mean difference = 137.4 normalized counts; P = 0.001)(Figure S6). The effect was not always consistent among genes, however, because 58.5% of allelic genes had lower expression in pcM alleles, while 41.4% had higher expression in these genes.

The second hypothesis is that pcMs correlate with temperature. We tested the hypothesis that the frequency of pcMs covaried with climate by assessing the frequencies of derived pcM alleles in geographically distinct subpopulations. To define subpopulations, we first used admixture groups determined from previous analyses of the 1001 Genomes (1001 Genomes Consortium 2016). For each subpopulation, we estimated the mean frequency of derived alleles. To compare population frequencies to climate data, we extracted climatic variables for each individual in each subpopulation based on its geographical coordinates and then calculated the mean of each climatic variable in each subpopulation. Several climatic variables related to temperature were negatively correlated with the frequency of syn_pcM alleles across subpopulations (Figure 5AB). The correlation was significant for BIO1 (mean annual temperature; linear model R_2_ = 0.51, P = 0.032), BIO6 (minimum temperature of the coldest month; R_2_ = 0.54, P = 0.024), and BIO11 (mean temperature of the coldest quarter; R_2_ = 0.47, P = 0.043). Other temperature-related variables and all precipitation-related variables not significantly correlated (P > 0.05; Figure S7). We repeated this analysis with syn_upM frequencies to test whether these results reflect the importance of secondary structure per se or geographic effects, finding that syn_upM regressions were significant for BIO1 (P = 0.047) but not for BIO6 (P = 0.052), BIO11 (P=0.062) or other bioclimatic variables.

**Figure 5.**
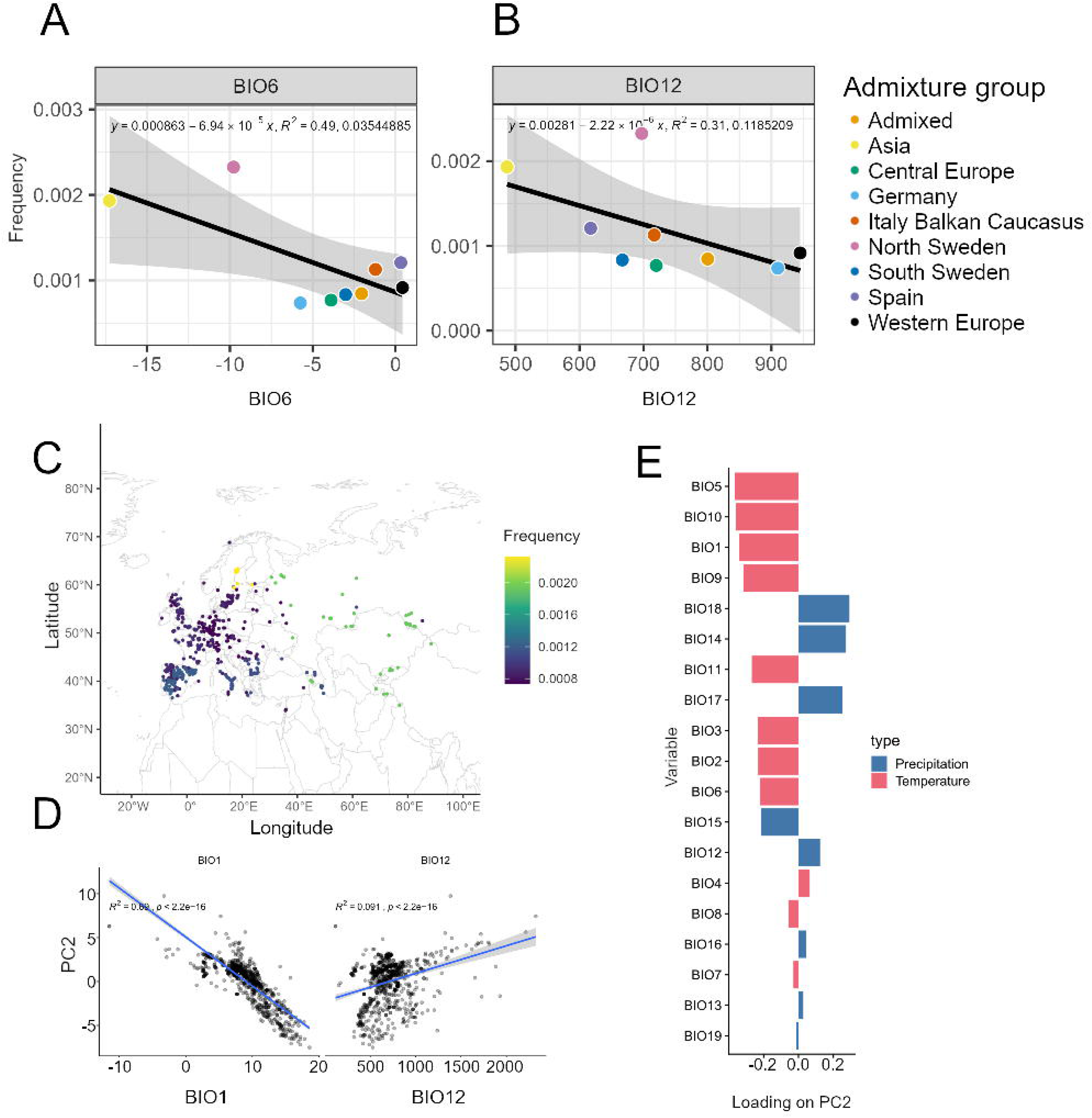
pcM associations with bioclimatic variables. A) The mean pcM frequencies within admixture groups as a function of the mean climate variables for BIO1, the Annual Mean Temperature. The equation shows the linear model, the inferred correlation coefficient (R_2_) and the p-value. B) As panel A, but for BIO12, Mean Annual Precipitation, which has a lower and non-significant relationship with allele frequency. BIO1 and BIO12 are shown as examples, with additional bioclimatic variables provided in Figure S7. (C) A map of Eurasia with the sampling location of the 1,001 Genomes, color-scaled by their syn_pcM allele frequencies. D) Linear models of PC 2 as a function of BIO1 and BIO12 variables for each individual. (E) Loading scores of environmental variables on PC 2. Greater values indicate that variables contribute more to PC 2, and variables are colored by class (temperature vs precipitation). Temperature-associated variables tend to have greater loading values than precipitation.

These results relied on the definition of populations based on admixture groups. Accordingly, we adopted an alternative approach that relied on individuals rather than previously-defined populations. We first summarized environmental variation across individuals using principal component analysis (PCA). The first three PCs explained 36.02%, 31.33%, and 12.93% of the variation in climate, respectively. We then included these first three PCs in mixed-linear models to predict the number of all pcMs, syn_pcMs and non_pcMs. We did not detect any significant associations between all pcMs and non_pcMs, but syn_pcMs were significantly associated with PC 2 (p = 0.00099). PC 2 also was geographically defined, with a strong correlation with latitude (Spearman’s ρ= 0.82, p = <2.2 ×10^-16^), and a moderate correlation with longitude (Spearman’s ρ = 0.34, p <2.2 × 10^-16^, Figure 5C). To understand the relative contribution of bioclimatic variables to PC 2, we examined the loadings of each bioclimatic variable on this axis. Bioclimatic variables related to temperature (BIO1 to BIO11) disproportionately contributed to PC 2 compared to precipitation-related bioclimatic variables; e.g., the absolute value of cumulative loadings across temperature variables was 2.48 on PC2 compared to 1.24 for precipitation variables (Figure 5DE). Supporting this observation, annual mean temperature (BIO 1) explained 69% of variation in PC 2 in a simple linear model, while annual precipitation (BIO 12) only explained ∼9% (Figure 5DE). Taken together, our analyses suggest that the distribution of pcMs is explained in part by differences in temperature across the Eurasian range of *A. thaliana*, but there is also evidence that these associations hold for upM mutations.

## DISCUSSION

Mutations that affect secondary structure have long been known to have functional effects (Wan et al. 2014). For example, they contribute prevalently to human genetic disease (Halvorsen et al. 2010; Lin et al. 2020) and a subset may act as riboSNitches that affect both secondary structure and gene expression (Ferrero-Serrano et al. 2022) in a temperature dependent manner (Su et al. 2018).

Here we have identified derived mutations that are likely to change pairing between bases within CDS regions, based on both bioinformatic predictions and experimental data. Of course, their identification is subject to numerous caveats. We relied, for example, on a pseudo-ancestral genome that was calculated from the Col-0 reference and an A. lyrata outgroup. This approach may mis-assign ancestral states for a subset of sites and excluded SNPs from the 1,001 genomes dataset that did not align reliably with A. lyrata. As a consequence, we almost certainly identified only a subset of pair-changing variants. Our analyses were also likely biased toward studying secondary structures that are present in the Col-0 reference, because we used dsRNA data from Col-0 in our identification pipeline.

To identify bases in coding regions that pair with other bases, we chose LinearPartition, because it performs reliably and efficiently relative to other secondary structure prediction programs (Zhang et al. 2020), including in extensive comparisons to RNAfold (Lorenz et al. 2011). We also used dsRNA data to verify pairing, even though Arabidopsis is one of only two plant species with available genome-wide structure-seq information (Ding et al. 2014; Deng et al. 2018). We investigated the use of structure-seq data, but most genes were not represented in the dataset, prohibitively limiting the possibility for genome-wide analyses. Finally, we assumed cutoff probabilities > 0.9 with LinearPartition to filter pair-changing mutations. Although the general results held with relaxed criteria, we suspect that our strict criteria misidentified many bona fide pcMs as either equivocal or unpaired mutations (upMs). The net effect of misclassifications is to underemphasize differences among site classes (pcM, equivocal and upM), making comparisons among categories inherently conservative.

### The case for selection against pair-changing mutations

Given our methods, we identified >200,000 bases across the expressed regions of the genome that likely pair with other bases on the same transcript. Only a subset of 8,469 bases were polymorphic across the Arabidopis 1,000 genomes dataset (Table 1), and these were the focus of our study. The examination of these pcMs suggest that they are under selective constraint, based on evidence that includes: i) an 13.4% reduction of nucleotide diversity (π) at syn_pcM sites compared to syn_upM sites (Figure 2A), ii) a skewed SFS in syn_pcM sites relative to syn_upM sites (Figure 3A), and iii) a strong underrepresentation of pcM changes in some genic locations, especially UTRs (Figure 2B). In addition to summary statistics, we inferred the DFE for various classes of sites based on a fitted demographic model. The DFEs reflect strong evidence for selection against derived non_upM variants (Figure 4A), as expected, but also for pcMs in UTRs (Table 3 & Figure 4B). As we discuss below, the case for selection against syn_pcM variants was less clear based on DFE analyses (Table 3) even though supported by other metrics.

The inference about purifying selection on derived pcMs within UTRs is consistent with previous work that has documented evolutionary constraints on secondary structure. Secondary structure has been shown to impose constraint on experimental evolution in microbial systems (Chursov et al. 2013; Bailey et al. 2021), and phylogenetic approaches have shown that evolutionary slower rates at synonymous sites correlate with the strength of secondary structure (Park et al. 2013). We have found that the 3’ regions of genes have marked reduction of nucleotide diversity for pair-changing mutations (Figure 2B), that UTRs in both 5’ and 3’ regions have skewed SFSs (Figure S4), and statistical support for selection based on DFE analyses of pcMs (but not upMs in UTRS) (Table 3). The remaining question is the functional basis for selection against derived pair-changing mutations. In Arabidopsis, it is known that the interruption of secondary structures in 3’ UTRs destabilize mRNAs (Zhang et al. 2024), and so it is likely that selection on mRNA half-lives or degradation rates could play a role. Similarly, 5 UTRs are generally tied to crucial functions in ribosome binding and translation (Babendure et al. 2006; Matoulkova et al. 2012). It is worth noting that secondary structures are common within the UTRs of plant genes; 85% of maize genes have detectable secondary structures in their 5’ UTRs (Martin et al., 2023) and rice and maize generally seem to have stronger folding dynamics in 5’ UTRs (Deng et al. 2018; Martin et al. 2023) compared to Arabidopsis (Deng et al. 2018). The abundances of genic transcript also vary with the strength of secondary structures. In maize, for example, transcript abundance is lower for genes with particularly strong and weak folding within their 5’ UTRs (Martin et al., 2023), suggesting that there are optimal folding parameters with respect to gene expression and translation. Altogether, selection against derived pcM mutations in UTRs may reflect their effects on gene expression, transcript stability and/or translation efficiency.

In contrast to UTRs, the case for selection against derived syn_pcMs is more circumspect, even though it has long been known that synonymous mutations are not entirely neutral (Ikemura 1981). For example, strongly deleterious synonymous variants have been documented in Drosophila, but the selective effects did not correlate with the strength of selection on secondary structure in this study system (Lawrie et al. 2013). Here most of our observations are consistent with the idea that derived syn_pcM sites are under slightly stronger negative selection than syn_upM sites, based on lower nucleotide diversity (Figure 2A), a skewed and significantly differently SFS (Figure 3A) and downstream associations with temperature and gene expression. The DFE analyses do not, however, necessarily support this conclusion. Relative to “neutral” syn_upM sites, the DFE analyses suggest that the syn_pcM variants are at most moderately deleterious, and their effects cannot be differentiated statistically from presumed neutrality (Table 3).

The syn_pcM DFE results likely reflect the some semblance of truth, in that syn_upM and syn_pcMs seems to be under fairly similar magnitudes of selection. However, the methods are also likely to lack discriminatory power, due to the fact (for example) that fitDadi and similar may not deal adequately with the effects of linked selection (e.g., Gilbert et al., 2022). Another limitation is that fitDadi was developed for the analysis of outcrossing species, but *A. thaliana* is predominantly selfing. Selfing can lead to decreased effective recombination rates, which in turn increases the potential for interference among linked alleles. Various ways have been implemented to deal with selfing in DFE analyses (Huber et al. 2018; Blischak et al. 2020), but empirical studies on the effect of selfing have been mixed. DFEs can be overestimated (i.e., inferring too much strong selection) when ∼100% selfing is not considered (Gilbert et al., 2022) but other studies have recovered adequate DFE distributions with selfing rates similar to Arabidopsis (Arunkumar et al., 2015; Huber et al., 2018). Another recent study has shown that inbreeding reduces the inferred selective effects of moderately deleterious alleles (Daigle and Johri, 2025), suggesting that our DFE-based estimates based on syn_pcMs may be conservative. We did attempt to use selfing models in fitDadi, but were unable to get them to converge with our data. Nonetheless, our demographic fits with outcrossing models were reasonable, and mean DFEs estimates were not obviously inflated for some categories of sites (e.g., syn_pcMs, UTR_upMs, etc.).

Our work has reinforced that genes have many potential targets of selection—from UTRs to missense changes to secondary structure—that could, in theory, lead to interference and complex trade-offs. For example, mutations in UTRs are likely to compete with other, linked changes in coding regions. There are also possible conflicts between the RNA versus protein life stages of a gene at individual sites; that is, mutations could be detrimental for secondary structure but advantageous for protein function, or vice versa. We assessed how often, at individual sites, the magnitude of negative selection against secondary structure (i.e., RNA level) changes was stronger than for nonsynonymous (i.e. protein level) changes, based on draws from the syn_pcM and non_upM DFE distributions. The results were surprising, because they showed that nearly half of genes may have at least one nonsynonymous mutation that has larger fitness effects due to effects on secondary structure compared to the encoded amino acid change. Thus, selection at RNA-level may often affect proteins. We recognize that our approach to investigate this phenomenon was simplistic, in that it assumed the inferred DFEs were accurate and also treated each gene equivalently with respect to both evolutionary rates and the probability of an amino acid change. Further disentangling the numerous (perhaps contradictory) pressures shaping gene evolution at both RNA and protein levels will require integrating structural dynamics into molecular and population genetic analysis.

### Paired mutations associate with temperature and gene expression

In a landmark study, Su et al (2018) presented a compelling demonstration of the potential importance of secondary structures within plant genes. They subjected rice (Oryza sativa L.) seedlings to a high temperature, eliciting a heat-shock response, and then found that the RNAs of ∼14,000 genes unfolded over the experimental temperature range. As a class, these genes also demonstrated shifts in gene expression, which they attributed primarily to more rapid degradation of unfolded RNAs (as opposed to reduced translation rates). Their work suggested that SNPs that modify secondary structures could be more or less tolerated, depending on temperature and climate. That is, selection against pcMs may not be as strong for plants that reside in regions of moderate (as opposed to high) temperatures. Their results also suggest that RNA folding is a vital component of gene expression, so that one expects correlations between the presence of secondary structure altering SNPs and shifts in gene expression.

To explore these threads on a genome-wide scale, we examined the distribution of pcM variants across the sampling landscape of the Arabidopsis 1001 Genomes dataset. We investigated the association of alleles and climate across subpopulations (i.e., previously inferred admixture groups) and across individuals. Both approaches provide genome-wide evidence that derived pcM mutations are more common at locations with lower temperatures, as measured by bioclimatic variables (Figure 5), without correspondingly strong associations to precipitation-based variables (Figure 5E). This is the first demonstration of this genome-wide pattern, to our knowledge, and it provides an opportunity to consider the evolutionary forces that contribute to such a pattern. We can think of three reasonable explanations: local adaptation, deleterious load and genetic/geographic clustering. Previous work has argued convincingly that some associations between pcMs and climate likely represent local adaptation events (Ferrero-Serrano and Assmann 2019), but we favor the latter two explanations for our genome-wide pattern, for two reasons. First, derived, deleterious pcM mutations may be less strongly selected against in low temperature environments where strong-folding may not be as critical, and deleterious mutations tend to accumulate at the edges of geographic ranges (Travis et al. 2007; Excoffier et al. 2009; Angert et al. 2020). Visually, we find that higher pcM counts occur in the Northern and Eastern edges of the sampled range (Figure 5C), perhaps representing expanding edges from Ice Age refugia. Second, syn_upM mutations also correlate with BIO1, suggesting associations among temperature, geography and genetic diversity.

Of course, any argument for selection for or against derived pcMs assumes that they have a phenotypic effect. We found the potential for such an effect, because genome-wide allelic expression was significantly lower for alleles with a derived pcM. This genome-wide result, across all sampled genes, mimics similar results in microbial systems where the disruption of secondary structures reduces gene expression (Bailey et al. 2021). However, there was also wide variation across genes, because >40% of genes showed the opposite pattern— i.e., pcM alleles had higher expression. We frankly find it surprising that we could detect any trend at all, given that alleles in most genes undoubtedly differ by more than just the presence/absence of a pcM and the concomitant experimental noise. The results suggest, although it is far from proven, the disruption of secondary structures has a causal effect on expression. One potential biological explanation for higher expression of pcM alleles is that the mutations that disrupt especially strong secondary structures may also interrupt RNA-interference (Li et al. 2012), thereby diminishing epigenetic control. No matter the cause, we have shown that derived pair-changing mutations are under moderate levels of purifying selection based on most of our analyses, that they vary across the genic location (e.g., UTR vs. synonymous sites), that they associate with temperature, and that one potential cause of these effects is that the perturbation of secondary structures alters the dynamics of transcript abundance.

## MATERIALS and METHODS

### Identification of derived upM and pcM mutations

We used the 1001 Genomes Project v.3.1 (https://1001genomes.org/data/GMI-MPI/releases/v3.1/) SNP calls (Lamesch et al. 2012) and variant annotations (1001 Genomes Consortium 2016) for all analyses. We filtered the variant dataset that included all 1001 genomes to retain biallelic SNPs and assigned ancestral and derived states by aligning the A. lyrata v1.0 genome assembly (Hu et al. 2011) to the *A. thaliana* TAIR10 reference (Lamesch et al. 2012) using AnchorWave 1.0 (Song et al. 2022). This approach enabled us to polarize 5,613,812 of 12,883,854 (43.6%) SNPs.

To identify sites that may contribute RNA secondary structures, we first constructed a pseudo-ancestral genome by replacing derived sites in the TAIR10 assembly with their corresponding ancestral alleles from the polarized VCF using GATK FastaAlternateReferenceMaker v3.7 (McKenna et al. 2010). Then, we extracted the longest mRNA (coding) sequence for each protein-coding gene from the pseudo-ancestral reference using bedtools2 getfasta 2.27.1 (Quinlan and Hall 2010) before estimating RNA folding for each sequence with LinearPartition v1.0 (Zhang et al. 2020). We then selected SNP sites that overlapped with positions with base-pairing probability > 0.9 as determined by LinearPartition for further analysis. We verified putative pairing sites by determining the overlap with dsRNA sequencing data generated from wildtype flower buds (NCBI Gene Expression Omnibus GSE23439) (Zheng et al. 2010). Since the dsRNA data was mapped to the TAIR9 assembly, we converted to TAIR10 assembly coordinates (Lamesch et al. 2012) using CrossMap 0.6.4 (Zhao et al. 2014). We considered putative pair changing mutations (pcM) with both computational and empirical evidence (i.e., high base pairing probability in LinearPartition analysis and dsRNA coverage). We further filtered these sites by finding overlap with potential compensating mutations using the base pairing probability files from LinearPartition; to do so, SNPs were excluded if the paired base position also contained an alternative allele with base-pairing compatibility with the derived allele. All overlaps of genomic features were calculated using the GenomicRanges R package 1.48.0 (Lawrence et al. 2013).

### Nucleotide diversity, allele frequency, and DFE analyses

We calculated nucleotide diversity (π) for pcM and upM sites using VCFtools v0.1.16 (Danecek et al. 2011). First, we extracted sites from the 1,001 genomes VCF file belonging to each category (syn_pcM, non_pcM, etc.) using samtools tabix, the 1,001 genomes SnpEff file and annotations from our paired/unpaired site identification. We calculated π with the per-site method in each gene using VCFtools. We measured distance between sites and various genic features (starts, stops, and intron junctions) using GenomicRanges in R.

Site frequency spectra were calculated using a custom R script with vcfR v1.15.0 (Knaus and Grünwald 2017), data.table v1.15.2 (Barrett et al. 2024), and tidyverse 2.0 (Kuhn and Wickham 2020) R packages. Permutation tests for differences between SFS were done by sampling the number of upM sites, building a SFS for each sample, calculating the difference between the sample SFS and the true upM SFS in each bin and then repeating this procedure for 10,000 iterations to generate a distribution of differences for each bin under the null model.

To estimate the DFE for different mutation types, we used the fitDadi python package (Gutenkunst et al. 2009; Kim et al. 2017). We first inferred demography using the syn_upM SFS, which we assumed represented neutral mutations, as in theory they affect neither the RNA secondary structure nor the protein product. We fit four demographic models (Table 3) by loading the syn_upM SFS into Fitdadi from the polarized VCF file and by projecting down to 50 frequency bins to moderate the effects of missing data. We optimized parameters for each demographic model using the multinomial method, and we perturbed each starting parameter at least 5 times to ensure that the same optimum for each demographic parameter was reached independent of starting values. We evaluated the accuracy of the inferred demographic models using fitDadi to generate a neutral SFS under each demographic model and compared the simulated SFS to the SFS from real data using Kolmogorov-Smirnov test in R. The estimated parameters for each model are provided in Table S1 and examples of model fits are provided in Figure S3.

We then used each of the fitted demographic models to estimate the DFE of syn_pcMs, UTR_pcMs, non_pcMs and non_upMs separately by using the unfolded SFS for each mutation type in fitDadi (Kim et al., 2017) modeling the DFE as a gamma distribution. We plotted DFEs in python using matplotlib (Hunter May-June 2007). We estimated the mean scaled selection coefficient of each gamma distribution by multiplying the shape × scale parameter of each. For both variant classes, we used likelihood ratio tests with two degrees of freedom to compare nested models that inferred the two gamma parameters (i.e., with and without the DFE) (Table 3). We also tested two separate syn_upM SFS for demographic inference: (1.) using a folded SFS and (2.) using subsampling instead of projection. We tested these alternative approaches because (1.) the unfolded SFS is dependent on accurate ancestral state-calls, and (2.) subsampling allows for modeling of inbreeding during optimization of the demographic model, while projection does not. However, we ultimately did not include these results because in both cases the model fits were much worse (growth model from folded SFS AIC = 756.66; subsampling growth model with inbreeding AIC = 528,690)

### Pleiotropy simulation

To investigate the potential for conflicts between protein and RNA-level selection, we started with a collection of 27,206 genes and multiplied the CDS lengths of each *A. thaliana* gene with an *A. lyrata* ortholog (downloaded from Ensembl) by 0.66 to approximate the number of nonsynonymous sites across the genome. For each gene, we assigned each site as either paired or unpaired based on the probability data from Table 1. We then assigned each site a “protein-level” fitness effect (γ) by pseudo-randomly drawing a value from the non_upM DFE gamma distribution (shape = 0.24, scale = 52.7). The assigned selection coefficient was pseudo-random because the maximum value of assignments was capped at 1,000 (2N_A_S). We then assigned paired sites an “RNA-level” fitness effect by the same method, but this time sampling from gamma distribution representing the DFE of synonymous pcM SNPs (shape = 0.22, scale = 73). For each simulation we counted the number of sites across the genome where selection was stronger (more negative) against secondary structure changes than amino acid changes. We evaluated the accuracy of our DFE simulations by comparing the means of these sampled DFE to the “true” means estimated from the gamma distributions (shape × scale). We repeated the simulation 100 times, finding that the results changed minimally (Figure 4).

### Geospatial and climatic correlations

We studied the association between environmental variables and the number of pcM SNPs at both subpopulation and individual scales. For the environmental variables, we used the 19 WorldClim 2 bioclimatic variables at 2.5 minute resolution, which summarize past climate averages from 1970-2000 (Fick and Hijmans 2017). Bioclimatic values for each accession were extracted using the collection coordinates reported by the 1001 Genomes Project (1001 Genomes Consortium 2016) and the raster 3.6-26 R package (Hijmans 2023). For the subpopulation-based approach, we considered the 10 subpopulations inferred previously (1001 Genomes Consortium 2016) and fit both simple linear models and generalized linear models in R (R Core Team 2023) to predict the mean allele frequency across syn_pcM alleles using the mean of each bioclimatic variable for each subpopulation.

For the individual-scale approach, we first did a principal component analysis of the bioclimatic variables using the prcomp function in R (R Core Team 2023) and then fit mixed linear models using the lmekin function from the coxme 2.2-20 R package (Therneau 2024) to test for association between between the first three PCs and the number of pcM alleles across pcM sites per accession. A centered relatedness matrix calculated from all biallelic SNPs using gemma 0.98.5 (Zhou and Stephens 2012) was included as a random effect in the models. We corrected the p-values using the Bonferroni method and assessed significance at α = 0.05.

### Expression analyses

Expression data in the form of log normalized counts was downloaded from the NCBI Gene Expression Omnibus (GSE80744) (Kawakatsu et al. 2016), which includes expression data for 24,175 genes in the 727 Salk accessions from the 1,001 Genomes dataset. pcM overlap with genes was determined using the GenomicRanges library in R (Lawrence et al., 2013). Allelic state for each accession was determined using the 1,001 Genomes VCF file. Genes with no pcM allele were excluded from the analysis, and only two allelic states were considered: whenever an accession contained one or more pcMs in the gene, it was considered a pcM allele, irrespective of whether the pcM was the same SNP between alleles (e.g., if an accession contained a pcM at one position within a gene, and another accession contained a pcM at a different position, both were put into the same category of “pcM alleles”). The mixed-effect linear model was analyzed using the R package lme4 (Bates et al. 2015), and included pcM allelic state as a fixed effect and gene identity as a random effect, expressed as:

## DATA AVAILABILITY

All of the data used in this study are freely available. Custom scripts can be found at github.com/GalenTMartin/structure_selection and extra files can be found at https://figshare.com/projects/RNA_structure_evolution/221926. The NCBI Gene Expression Omnibus numbers for dsRNA and gene expressions were GSE23439 and GSE80744.

## ACKNOWLEDGEMENTS

We thank Edwin Solares for computing infrastructure support, as well as Kirk Lohmeuller and Chris Kyriazis for advice on demographic inference. This work was supported by NSF grants IOS-2414478 and IOS-2414478 to BSG.

