## Supplemental Files for "Quantifying the evolution of SNPs that affect RNA secondary structure in *Arabidopsis thaliana* genes"

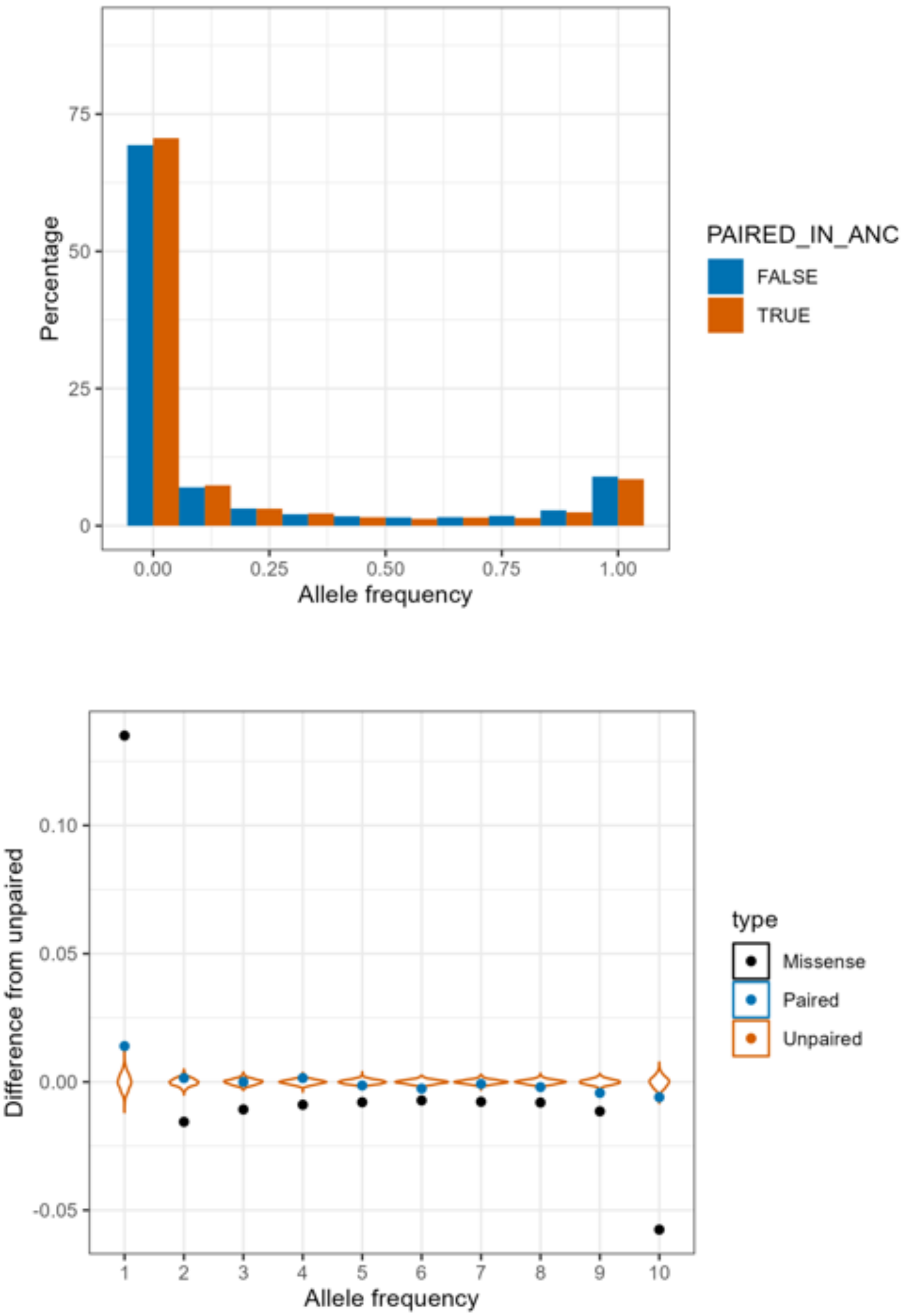


**Figure S1.** Top Panel) Site frequency spectrum and permutation tests for frequencies of synonymous pcM (True) and upM (False) defined without the requirement of dsRNA coverage (all sites with base pairing probability > 0.90 were considered paired sites). No equivocal category is included because only one metric of pairing was used (LinearPartition). Bottom Panel) Black dots represent differences between missense, as a point of reference, and synonymous paired (syn_pcM) and unpaired (syn_upM) mutation frequencies.

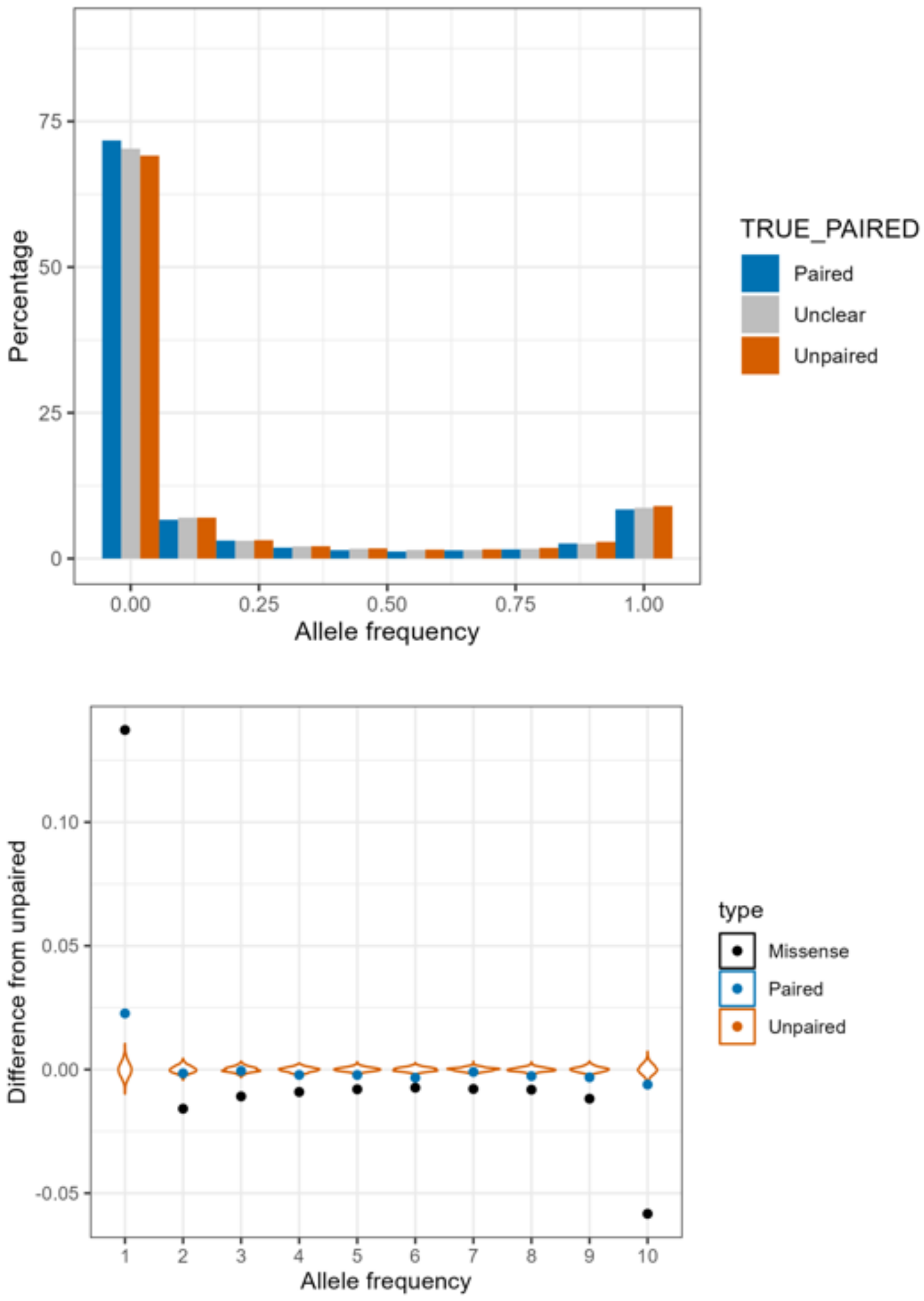


**Figure S2.** Top Panel) Site frequency spectrum paired (syn_pcM), equivocal (unclear), and synonymous unpaired (syn_upM) defined with a base pairing > 0.50 (50%) from linear partitions and also considering dsRNA coverage. Bottom Panel) Black dots represent differences between missense, as a point of reference, and synonymous paired (syn_pcM) and unpaired (syn_upM) mutation frequencies.

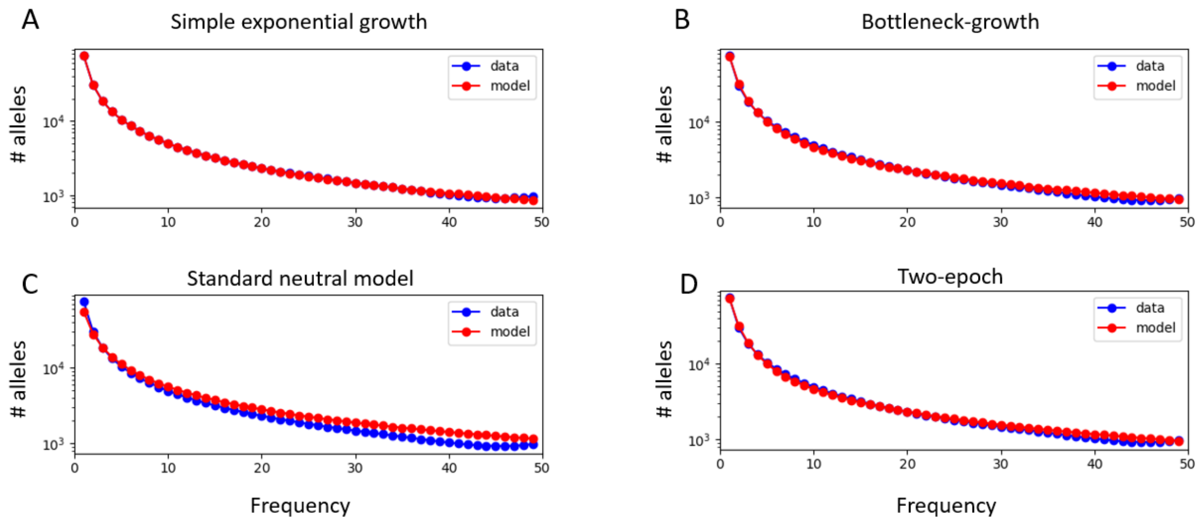


**Figure S3.** Site frequency spectra comparisons between the SFS estimated by fitDadi from optimized parameters in each demographic model (model, in red) versus the observed syn_upM SFS (data, in blue). (A) Simple exponential growth model, with growth from an ancestral population size. This was the best fitting model based in our analyses, based on AIC (Table 3). (B) Bottleneck growth, with a population reduction followed by exponential growth to contemporary population size, (C) a standard neutral model with constant population size, and (D) a two-epoch model with instantaneous population size change.

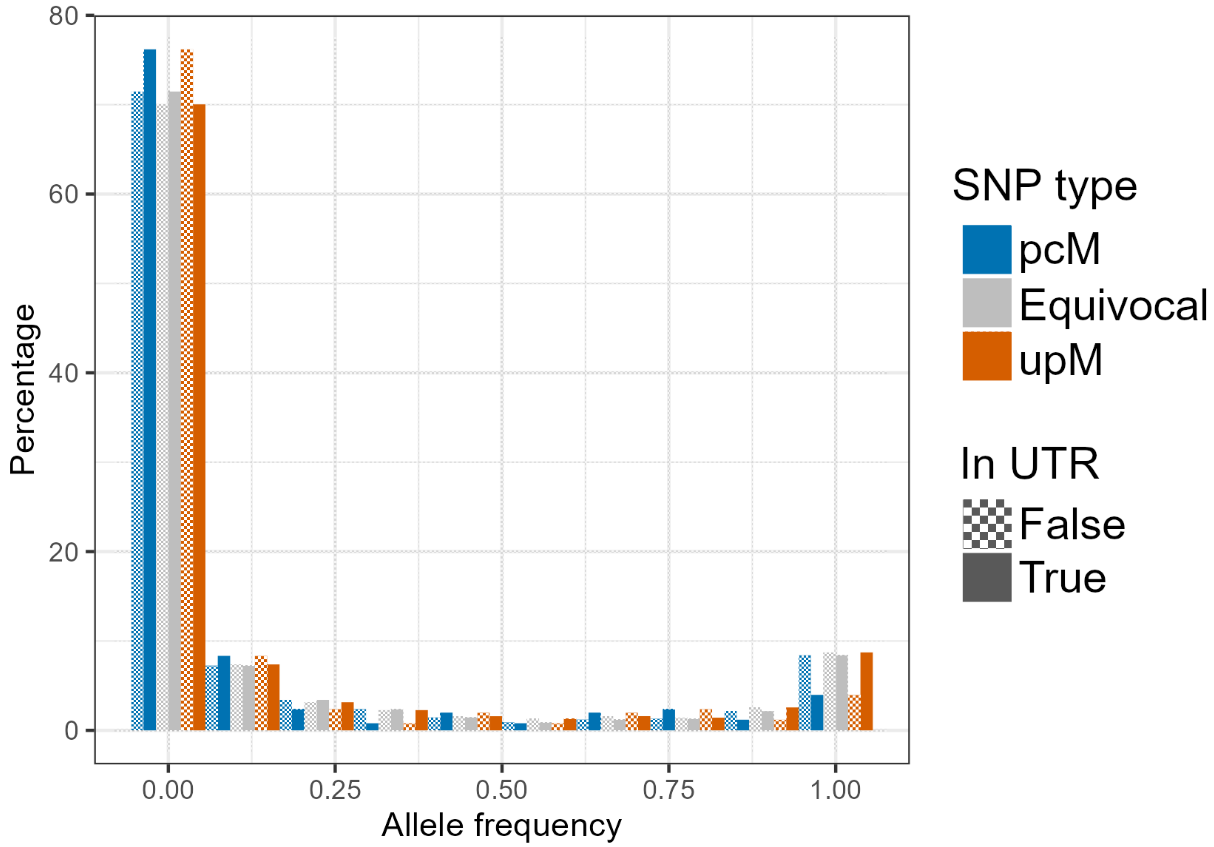


**Figure S4.** Site frequency spectrum for frequencies of pcM, equivocal, and upM within and outside of untranslated regions. Checkerboard-patterned bars show frequencies of alleles that are not in UTRs, while solid bars show alleles inside UTRs.

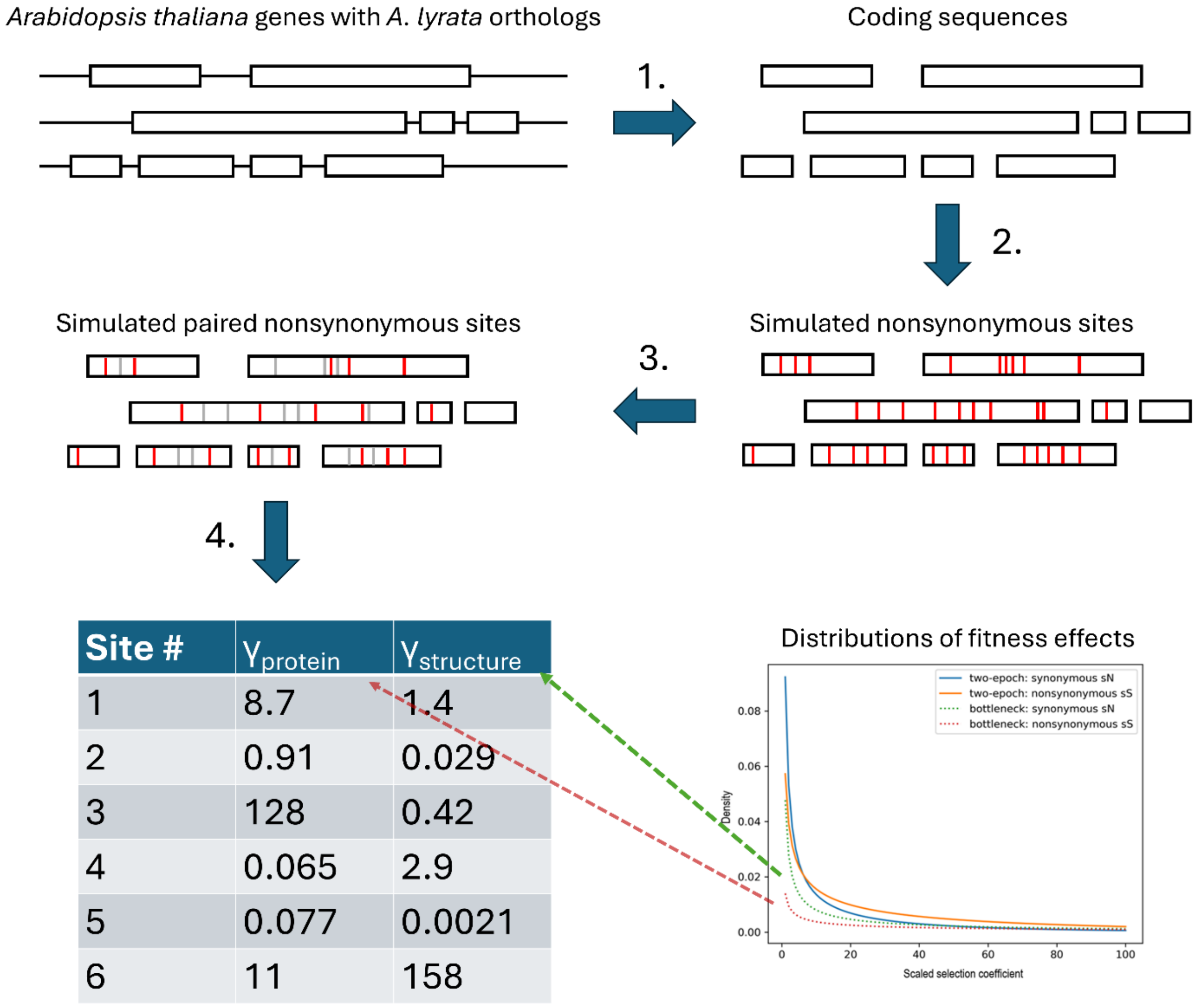


| Site # | $\gamma_{protein}$ | $\gamma_{structure}$ |
|---|---|---|
| 1 | 8.7 | 1.4 |
| 2 | 0.91 | 0.029 |
| 3 | 128 | 0.42 |
| 4 | 0.065 | 2.9 |
| 5 | 0.077 | 0.0021 |
| 6 | 11 | 158 |

**Figure S5.** Schematic of pleiotropy simulations. The simulations consisted of four steps as follows: (1). Extracted CDS lengths from Arabidopsis gene annotation, denoted by black boxes. (2). Assigned nonsynonymous mutations for each site based on frequency from Kimura (1968), denoted by red bars. (3). Assigned structure status of each nonsynonymous site based on our pairing frequency estimates (grey bars represent unpaired nonsynonymous SNPs, which were not considered further). (4). Assigned paired nonsynonymous sites two γ values based on estimated distributions of fitness effects. After the assignment, the number of sites with higher γ value for structure were counted, as were the number of genes that contained those sites.

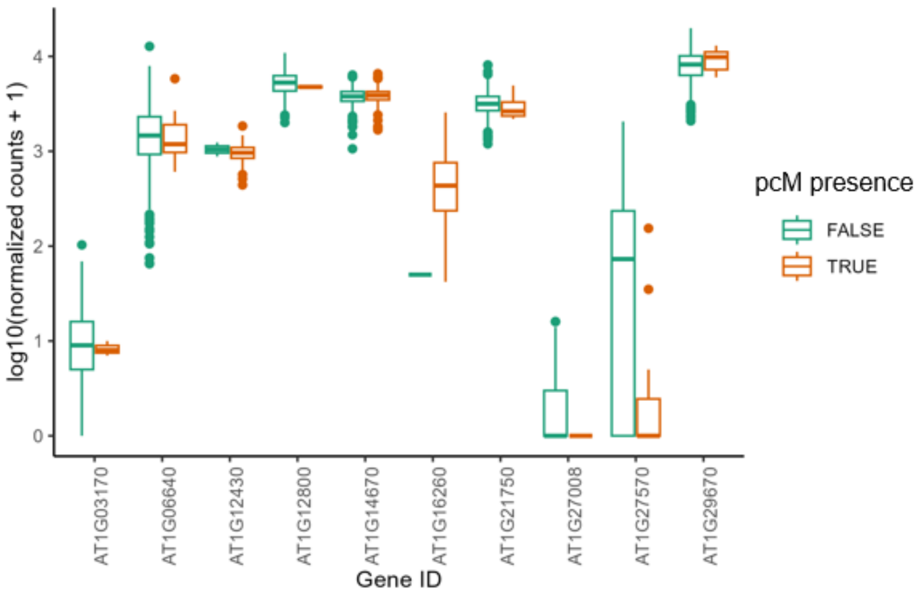


**Figure S6.** Expression between alleles with (TRUE) and without (FALSE) pcM mutations to illustrate the results for a handful of genes. The ten genes represent a random sample of genes from our analysis; they are meant to show that the mixed-effect linear model compared intra-gene expression differences between allelic states, that the differences between pcM and upM alleles was often quite minor.  Nonetheless, the effect was significant genome-wide, with a documentable trend toward lower expression for pcM containing alleles.

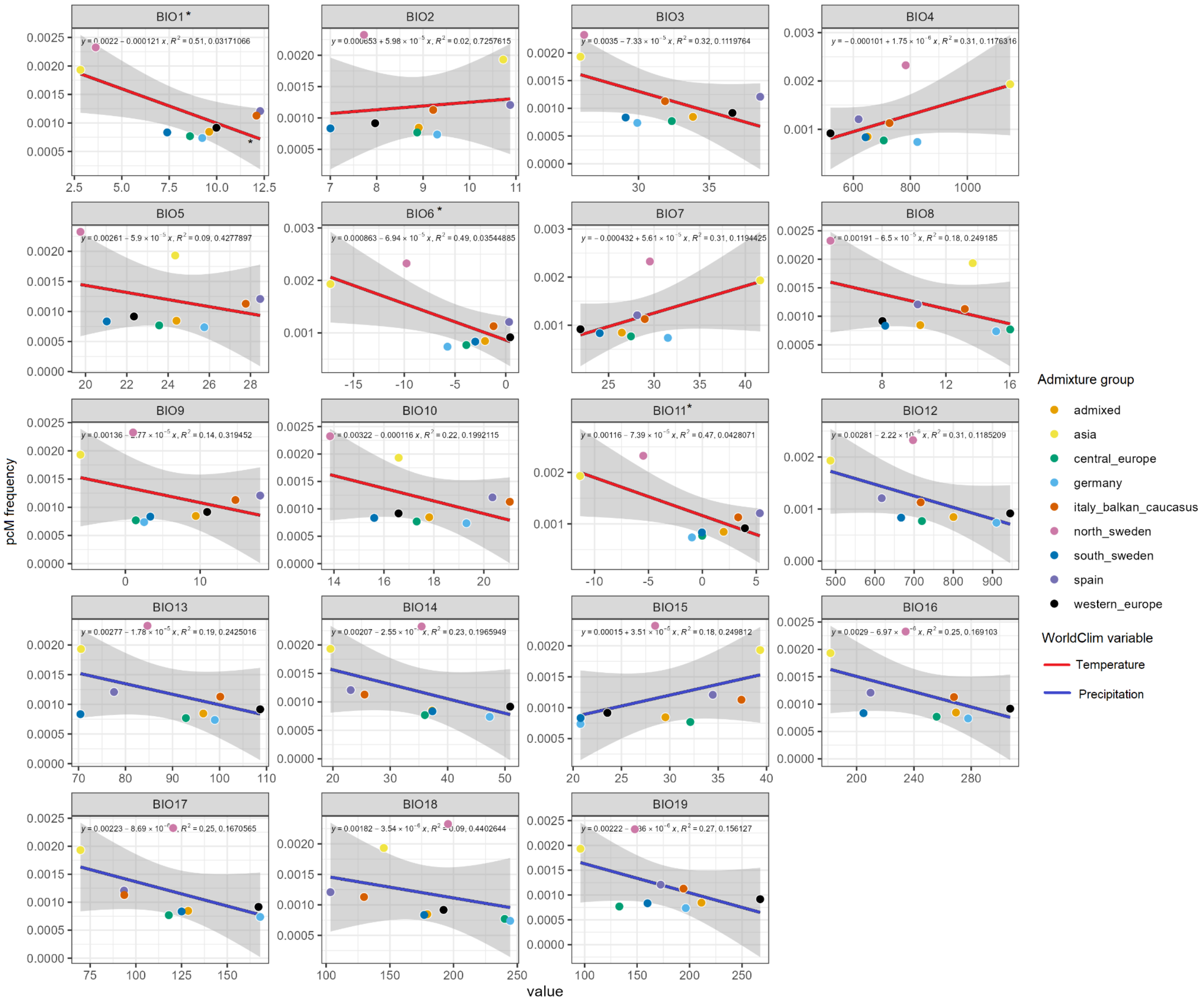


**Figure S7.** Linear regression between mean pcM frequencies in admixture groups and WorldClim variables. In each panel, lines are colored by variable category (temperature or precipitation), and asterisks show significant correlations ($P < 0.05$). Correlations were significant in BIO1 (mean annual temperature), BIO6 (minimum temperature of the coldest month), and BIO11 (mean temperature of the coldest quarter).

Table S1. Estimated demographic parameters

| **Demographic model** | **Parameters** | **nu**[1] | **T**[2] **($2N_{ancestral}$ Generations)** | **nuB**[3] | **nuF**[4] |
|---|---|---|---|---|---|
| Standard neutral model | - | - | - | - | - |
| Growth | nu,T | 2.26 | 0.243 | - | - |
| Two-epoch | nu,T | 2.03 | 0.129 | - | - |
| Bottleneck-growth | nuB,nuF,T | - | 0.225 | 0.98 | 2.26 |

1. nu = ratio of contemporary to ancestral population size

2. T = Time since population size change

3. Ratio of bottleneck population size to ancient population size

4. Ratio of population size after bottleneck recovery to ancient population size
